# The imprinted *Mir483* is a growth suppressor and metabolic regulator functioning through IGF1

**DOI:** 10.1101/2022.09.09.507324

**Authors:** Ionel Sandovici, Denise S. Fernandez-Twinn, Niamh Campbell, Wendy N. Cooper, Yoichi Sekita, Ilona Zvetkova, David Ferland-McCollough, Haydn M. Prosser, Lila M. Oyama, Danilo Cimadomo, Karina Barbosa de Queiroz, Cecilia S.K. Cheuk, Nicola M. Smith, Richard G. Kay, Katharina Hoelle, Noel H. Smith, Stefan H. Geyer, Lukas F. Reissig, Wolfgang J. Weninger, Kenneth Siddle, Anne E. Willis, Martin Bushell, Susan E. Ozanne, Miguel Constância

**Affiliations:** Metabolic Research Laboratories and MRC Metabolic Diseases Unit, Wellcome Trust-MRC Institute of Metabolic Science, University of Cambridge, Cambridge, United Kingdom; Department of Obstetrics and Gynaecology and National Institute for Health Research Cambridge Biomedical Research Centre, Cambridge, United Kingdom; Centre for Trophoblast Research, Department of Physiology, Development and Neuroscience, University of Cambridge, Cambridge, United Kingdom; Present address: Cancer Research UK Cambridge Centre, Cancer Research UK Cambridge Institute, Li Ka Shing Centre, Cambridge, United Kingdom; Present address: Laboratory of Stem Cell Biology, Department of Biosciences, Kitasato University School of Science, Kanagawa, Japan; Medical Research Council Toxicology Unit, University of Leicester, Leicester, United Kingdom; Present address: Institut de Recherches Cliniques de Montréal, Montreal, Quebec, Canada; The Wellcome Trust Sanger Institute, Genome Campus, Hinxton, United Kingdom; Present address: Cambridge Institute of Therapeutic Immunology and Infectious Disease, and Department of Medicine, Jeffrey Cheah Biomedical Centre, University of Cambridge, Cambridge, United Kingdom; Departmento de Fisiologia, Universidade Federal de São Paulo, Escola Paulista de Medicina, São Paulo, Brasil; Laboratory of Developmental Biology, Department of Biology and Biotechnology “Lazzaro Spallanzani”, University of Pavia, Pavia, Italy; Present address: Clinica Valle Giulia, GeneraLife IVF, Rome, Italy; Present address: Departamento de Alimentos, Programa de Pós-Graduação em Saúde e Nutrição, Escola de Nutrição, Universidade Federal de Ouro Preto, Ouro Preto, Brasil; Nuffield Department of Women’s & Reproductive Health, University of Oxford, Oxford, United Kingdom; Present address: Norwegian Institute of Public Health, Oslo, Norway; Lonza Biologics, Chesterford Research Park, Saffron Walden, United Kingdom; Center for Anatomy and Cell Biology, Division of Anatomy, Medical University of Vienna, Vienna, Austria; Present address: Medical Research Council Toxicology Unit, University of Cambridge, Cambridge, United Kingdom; Present address: Cancer Research UK Beatson Institute, Glasgow, United Kingdom and Institute of Cancer Sciences, University of Glasgow, Glasgow, United Kingdom

## Abstract

*Mir483* is a conserved and highly expressed microRNA in placental mammals, embedded within the *Igf2* gene. Here, we uncover the control mechanisms and physiological functions of *Mir483 in vivo*, by generating constitutive loss-of-function and over-expressing mice. *Mir483* expression is imprinted and dependent on the *Igf2* promoters and *Igf2/H19* imprinting control region. Over-expression of *Mir483* causes severe mid-gestation fetal, but not placental, growth restriction, and late lethality. Fetal death is prevented by restoring *Mir483* to endogenous levels using an inducible transgenic system. Continuous postnatal *Mir483* over-expression induces growth stunting, elevated hepatic lipid content, increased adiposity, reduced local and systemic IGF1 levels and increased GH. The growth phenotypes are rescued by IGF1 infusion. Our findings provide evidence for a novel functional antagonism between a growth-suppressor microRNA and its growth-promoter host gene, and suggest that *Mir483* evolved to limit excessive tissue growth through repression of IGF ligand signalling.

## Main

MicroRNAs (miRs) are endogenous non-coding small RNAs that modulate gene expression at the post-transcriptional level and are critically involved in many cellular processes^1,2^. In most cases, miRs interact with the 3⍰UTR of target mRNAs to suppress their expression; however, miRs can also exert their action through binding at the 5⍰UTR or coding sequences of target mRNAs^3^. Aberrant expression of miRs is associated with a number of diseases, in particular various cancers. For this reason, miRs are increasingly used as potential biomarkers of disease^4,5^ and therapeutic agents^6^.

Imprinted domains, which are chromosomal regions containing clusters of genes expressed preferentially from one parental allele, transcribe hundreds of small non-coding RNAs, including small nucleolar RNAs (snoRNAs) and miRs^7^. It is estimated that ^~^7% of known human miRs are encoded by imprinted domains^8^. Most imprinted miRs are generated from three evolutionary different chromosomal domains, CM19MC (primate-specific), C14MC (eutherian-specific) and C2MC (rodent-specific), as large repetitive arrays^9,10^. They show tissue-specific expression, with marked or exclusive expression in placenta^11^, and with multiple placental roles (e.g. trophoblast invasion, migration, proliferation, differentiation, antiviral defence). However, there are a few imprinted miRs produced by a single gene locus^9,11^, including the imprinted *IGF2/H19* domain. This domain is part of the so-called ‘imprinting growth’ chromosomal region, on human 11p15 and distal mouse chromosome 7. Imprinted genes residing in these clusters have key functions in placental development and fetal growth. De-regulation of imprinted expression of a subset of these genes are causative of two congenital human growth syndromes (BWS – Beckwith-Wiedemann Syndrome and SRS – Silver-Russell Syndrome^12^), and a variety of cancers also show altered expression linked to tumour growth^13^.

The *Igf2/H19* domain contains two isolated miRs that are highly expressed in placental and fetal tissues: *Mir483* (mouse nomenclature^14^), which is embedded within an intron of the *Igf2* gene and *Mir675*, which is located in exon 1 of the *H19* gene (Fig. 1a). The paternal specific expression of *Igf2* and the maternal specific expression of *H19* is mainly under the control of a differentially DNA methylated region located upstream of *H19*, called Imprinting Control Region 1 (ICR1 or IC1)^7,15^. Work in mice strongly suggests that the controlled release of *Mir675* from the *H19* gene is important to limit the growth of the placenta specifically in late gestation^16^. The physiological roles of *Mir483* in a developmental context are unknown.

**Fig. 1:**
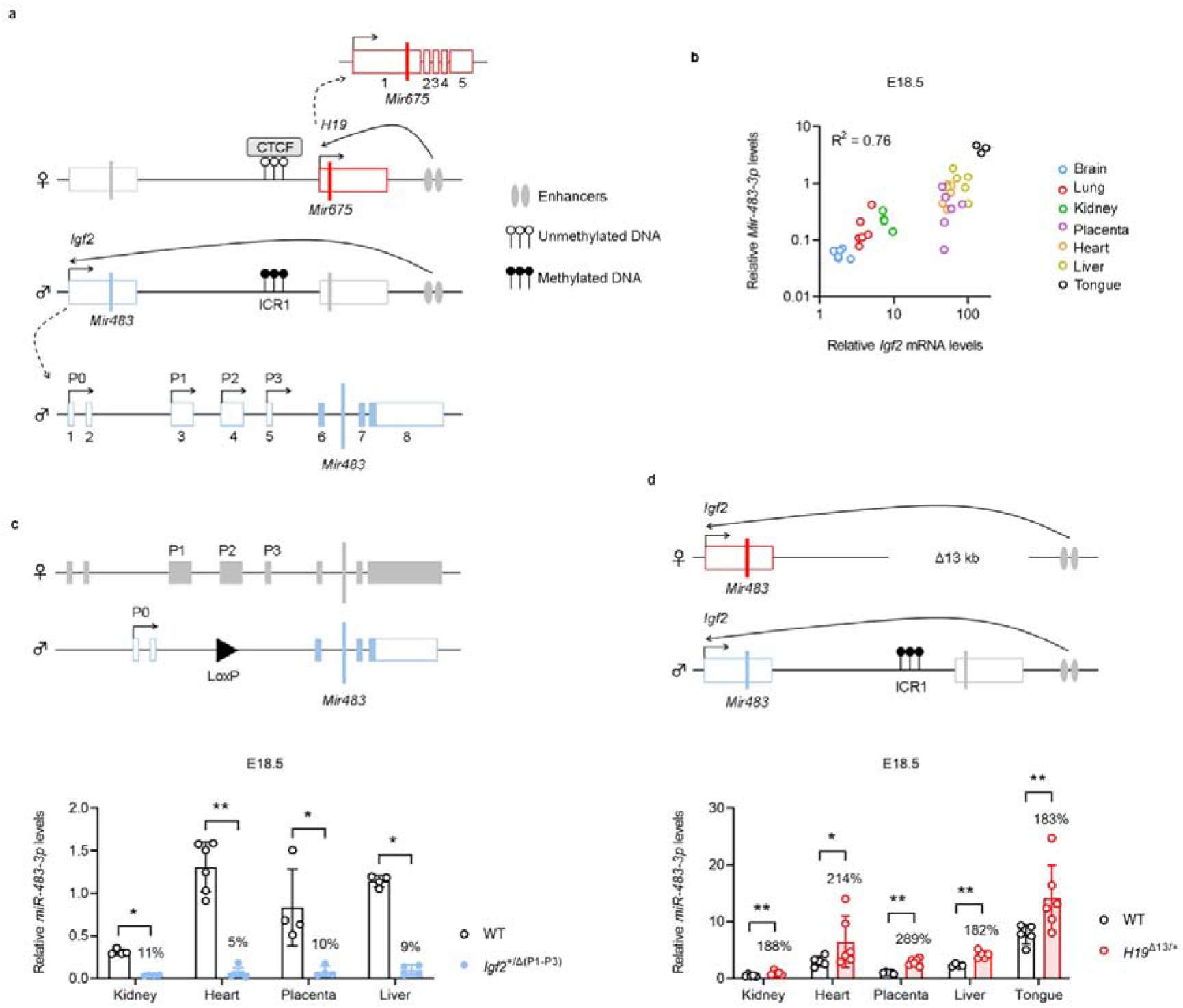
Developmental regulation of the mouse *Mir483* expression. **a,** Location of *Mir483* and *Mir675* within the *Igf2/H19* imprinted domain and regulation of *Igf2* and *H19* imprinting by the methylation-sensitive CTCF boundary at ICR1. P0 – P3 are alternative *Igf2* promoters and numbers indicate exons. **b**, Expression of *miR-483-3p* shows a strong positive correlation with *Igf2* expression in multiple fetal organs (two-tailed *P* value < 0.001; n=38 pairs with n=3-6 replicates per organ). Levels of *miR-483-3p* were normalized against *Snord70/snoRNA234*, and relative levels of *Igf2* were normalized against the geometrical mean of *Ppia, Pmm1* and *Hprt*. **c,** Deletion of fetal *Igf2* promoters P1-P3 from the paternal allele (*Igf2*^+/Δ(P1-P3)^ model), which are replaced by a loxP site, abolishes *miR-483-3p* expression at E18.5 (expression levels normalized against *Snord70/snoRNA234*). **d,** *Igf2* loss-of-imprinting (achieved by deletion of ICR1 and *H19* on the maternal allele – *H19*^Δ13/+^ model) results in increased relative levels of *miR-483-3p* expression in multiple fetal organs at E18.5 (normalized against *Snord70/snoRNA234*). For panels c and d, data are presented as individual values with averages ± standard deviation (SD); % values above the mutant columns indicate ratios mutant/wild-type (WT); n=4-6 samples per group; * P<0.05, ** P<0.01 by Mann-Whitney tests followed by two-stage step-up (Benjamini, Krieger, and Yekutieli) FDR < 5%.

Most of the published studies regarding *Mir483* are related to cancer, with the majority of the reports classifying *MIR483* (human nomenclature^14^) as an onco-miR^17,18^, though some studies provide evidence for an onco-suppressor action in certain contexts^19,20^. *Mir483* has been associated with a number of biological processes, such as cellular differentiation^21^, proliferation^22^, survival^23^, melatonin synthesis^24^, insulin production^25,26^, and vascular homeostasis^27^. Importantly, *Mir483* has been shown to be regulated by environmental cues such as diet^28^ and temperature^29^.

In this study, we investigated the mechanisms that regulate the expression and imprinting of *Mir483* and addressed functional roles during *in utero* and post-natal development using loss-of-function and gain-of-function *Mir483* transgenic mouse models. Our data suggest that a main physiological role for the *Igf2* encoded *Mir483* is growth control, mediated by the regulation of IGF1 levels.

## Results

### The mouse *Mir483* is under the regulatory control of *Igf2* and ICR1, and is not a self-regulating miR

*Mir483* is embedded within the intron 6 of the *Igf2* gene (Fig. 1a). The primary sequence and predicted *mir-483* (nomenclature for stem-loop^14^) secondary structure are highly conserved in eutherian mammals and marsupials, but not in monotremes (Extended Data Fig.1). The expression of mouse *miR-483-3p* (nomenclature for mature miR^14^) strongly positively correlated with that of the host *Igf2* gene across a range of fetal tissues (Fig. 1b). Moreover, both *Igf2* and *miR-483-3p* levels decreased in postnatal life, with very low expression observed in adult liver (Extended Data Fig. 2a) and other adult organs (Extended Data Fig. 2b). Promoter-specific transcript analysis showed that Promoter 2 (P2) was the *Igf2* promoter whose activity had the strongest correlation with *miR-483-3p* levels, in a range of tissues and across developmental time points (Extended Data Fig. 2b).

The expression associations between the *Igf2* and *Mir483* transcripts suggested that *Mir483* shares regulatory elements with the host gene. To test this hypothesis, we first generated and analysed mice carrying a deletion of the *Igf2* upstream transcriptional unit, *i.e*. main promoters P1, P2, P3 and the associated 5’UTR exons (referred to as *Igf2*^+/Δ(P1-P3)^) (Extended Data Fig. 3). We observed that *miR-483-3p* transcription was almost completely abolished in mice carrying this deletion on the paternal allele (Fig. 1c), in tandem with loss of *Igf2* expression (Extended Data Fig. 4a). Next, we investigated if the *Igf2/H19* imprinting control region – or ICR1 – located upstream of the *H19* gene (Fig. 1a) also controlled the expression of *Mir483*. A maternally inherited deletion of the ICR1/H19 gene (referred to as H19^Δ13/+^ – top panel in Fig. 1d), an established *in vivo* model of *Igf2* loss-of-imprinting in offspring^30^, led to the re-activation of the maternally silent *Igf2* promoters (Extended Data Fig. 4b). *MiR-483* levels were increased in parallel (Fig. 1d), at similar relative levels to those observed for *Igf2* (Extended Data Fig. 4b).

Expression of the human *MIR483* has been reported to be driven by an upstream miR-specific promoter^31^. Analysis of the human *MIR483* promoter sequence against the mouse showed little evidence of similarity, thus arguing against *Mir483* functioning as a self-regulating miR in the mouse (Extended Data Fig. 5a). We next tested for conservation in mice of the previously reported *miR-483-3p* seed matches in the human *IGF2* locus^18^. *In silico* analysis revealed putative sites for *miR-483-3p* regulation at the 3’ and 5’ mouse *Igf2* UTRs, with a binding site mapping to the mouse P2 promoter region (exon 4 5’UTR), equivalent to human P3 promoter region (exon 6 5’UTR), which was conserved in eutherians mammals and the marsupial wallaby (Extended Data Fig. 5b).

### Constitutive *Mir483* specific knockout does not show growth or metabolic phenotypes

Embryonic stem cells carrying a Cre-mediated deletion of *Mir483* were used to generate a *Mir483* knockout mouse (referred to as *Mir483*^Pat-KO^; see Fig. 2a and Extended Data Fig. 6). Paternal transmission of the deletion did not alter *Igf2* levels (Fig. 2b and Extended Data Fig. 7a,b), but caused >95% reduction in levels of *miR-483-3p* (Fig. 2b), providing unequivocal demonstration that *Mir483* is tightly regulated by genomic imprinting. There was no impact observed on the Mendelian distribution (n=21 WT and n=16 *Mir483*^Pat-KO^ in four mixed litters at E18.5, Fisher’s exact test p=0.65) or fetal and placental growth kinetics (Fig. 2c and Extended Data Fig. 7c). Postnatal growth of *Mir483*^PAT-KO^ mutants was indistinguishable from wild-type littermates (Fig. 2d,e and Extended Data Fig. 7d), as were fat mass, lean mass, bone mass density (Fig. 2f) and glucose tolerance (Fig. 2g).

**Fig. 2:**
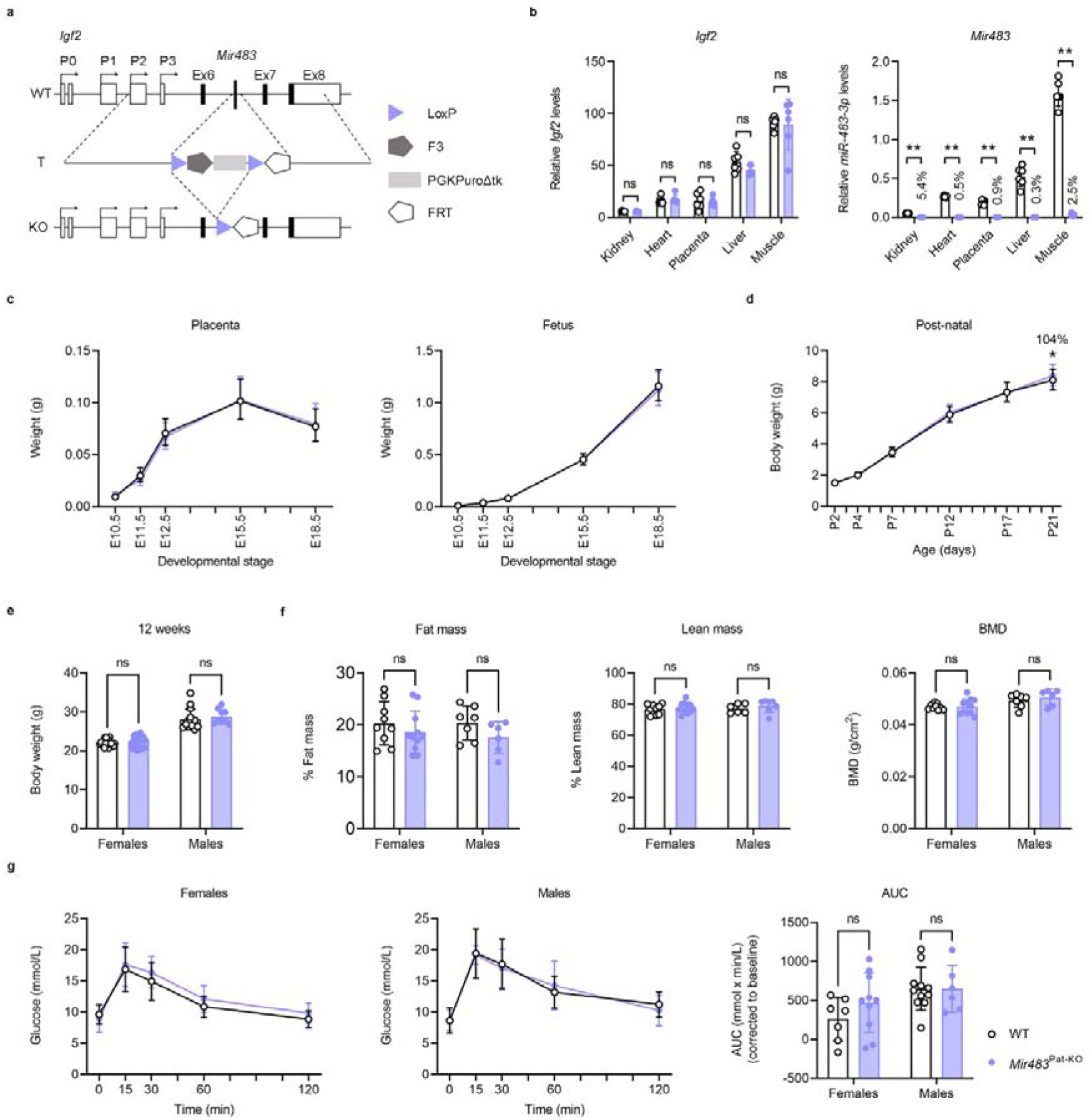
Developmental phenotyping of the *Mir483* knockout. **a,** Schematic representation of the *Igf2* wild-type allele (WT), targeting vector (T, containing the 5’ complementarity arm, loxP site followed by F3 site, PGKPuroΔTk selection cassette, a second LoxP site followed by a FRT site, and the 3’ complementarity arm) and the knockout allele (KO, obtained upon Cre-mediated deletion of the selection cassette). Details about the generation of the *Mir483* knockout are shown in Extended data Fig. 6. b, RT-qPCR levels of *Igf2* and *miR-483-3p* in E18.5 fetal organs from the knockout, upon paternal transmission of the deletion (*Mir483*^Pat-KO^), and wild-type (WT) littermate controls (n=6 samples per group). Levels of *Igf2* were normalized against the geometrical mean of *Ppia, Pmm1* and *Hprt; miR-483-3p* levels were normalized against the geometrical mean of *Snord70/snoRNA234* and *Snord68/snoRNA202*. **c,** Fetal and placental growth kinetics (n=4-6 litters at each developmental stage). **d**, Post-natal growth kinetics from post-partum day 2 (P2), until weaning (P21) (n=11-20 litters at each developmental stage). **e,** Body weights in 12 weeks-old knockouts and wild-type littermates (n=9-31 per group). **f**, Body composition (% fat mass, % lean mass and bone mineral density – BMD) measured by dual energy x-ray absorptiometry (DEXA) at the age of 12 weeks (n=6-11 per group). **g**, Glucose tolerance tests with glucose administered by intra-peritoneal injections (ipGTTs) after overnight fasting in females (n=7-10/genotype) and males (n=6-11/genotype). First two panels show changes in blood glucose concentrations (y-axis), from basal pre-treatment values, with time (x-axis), after glucose administration. The graph on the far right shows area under curve (AUC) calculated during ipGTTs using the trapezoid rule and normalised to basal glucose levels. Data are presented as individual values, with averages ± SD in **b, e, f** and **g** (far-right graph), averages ± 95% confidence intervals (95%CI) in **c** and **d,** and average ± SD in **g** (first two graphs on the left) and % values indicate ratios *Mir483*^Pat-KO^/WT; ns – non-significant, * P<0.05, ** P<0.01 by multiple Mann-Whitney tests with FDR (<5%) correction in **b**, a mixed effects model in **c** and **d**, and two-way ANOVA followed by Šídák’s multiple comparisons tests in **e,** f and g (far-right graph).

### Over-expressing *Mir483* causes fetal growth restriction and mid-gestation lethality

To fully establish the function of *Mir483* gain-of-function *in vivo* models were generated. In our initial approach, using homologous recombination in ES cells, we re-inserted one copy, three-tandem extra-copies and five-tandem extra copies of *Mir483* at the endogenous *Igf2* locus (Fig. 3a and Extended Data Fig. 8), but only achieved germ-line transmission of the five-tandem copy transgene (referred to as *Mir483*^5C^). Chimeric males transmitted the transgene to offspring, but the elevated levels of *Mir483^5C^* (observed for both *miR-483-3p* and *miR-483-5p*, see Fig. 3b) caused all the embryos to arrest in development, with complete reabsorptions by E13.5. At E11.5, fetuses, but not placentae, showed severe growth restriction (^~^58% of normal) (Fig. 3c) despite similar levels of *miR-483* overexpression in both the embryo and placenta (Extended Data Fig. 9a). Mass spectrometry-based protein quantification of ^~^4,000 proteins revealed a small number of significant changes (fold change > 1.5, with FDR <0.05) in whole embryos, related to induction of cytolysis (e.g. granzymes) and down-regulation of proteins implicated in epigenetic processes (e.g. histones) (Fig. 3d). The list of proteins down-regulated in *Mir483*^5C^ mutants included IGF2 (Fig. 3d and Supplementary Table 1). Down-regulation of IGF2 in *Mir483*^5C^ embryos, but not placentae, was validated by RT-qPCR and western blots in independent samples at E11.5 (Fig. 3e). Notably, the greatest mRNA reduction was observed for the *Igf2-P2* transcript (Extended Data Fig. 9b), which contains, as mentioned before, an 8-mer seed sequence in the 5’UTR exon 4 (Extended Data Fig. 9b).

**Fig. 3:**
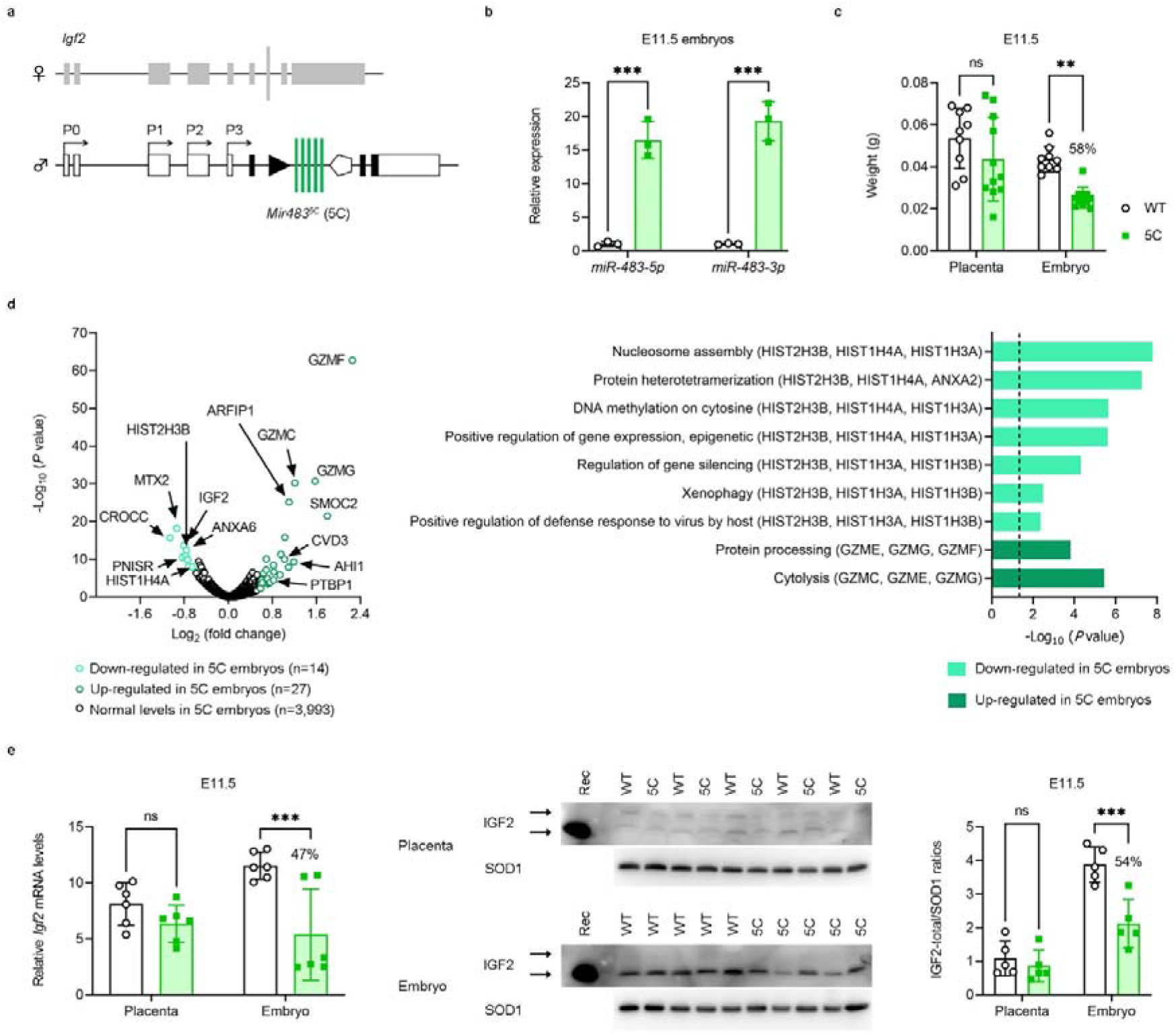
Overexpression of *Mir483* (*Mir483*^5C^ – 5C) from the endogenous *Igf2/Mir483* locus. **a,** Gene targeting of the 5-copy tandem array inserted at the endogenous locus (details about the generation of the *Mir483*^5C^ model are shown in Extended data Fig. 8). b, Relative expression of *miR-483-5p* and *miR-483-3p* measured by RT-qPCR in whole embryo lysates at E11.5 (n=3 samples/group). Levels of *miR-483-5p* and *miR-483-3p* were normalized against *Snord68/miRNA234*, and are presented relative to the wild-type (WT) levels, arbitrarily set to 1. **c,** Placenta and embryo weights at E11.5 (n=9-11/group). **d**, Proteomic analysis by TMT (Tandem Mass Tag) in surviving embryos at E10.5 (n=3 per genotype). Left: volcano plot representing all proteins quantified by TMT (see also Supplementary Table 1). Proteins that are significantly down-regulated or up-regulated (fold change >1.5, Benjamini-corrected P<0.00144) are presented in light-green and dark-green, respectively. Right: top scoring biological processes enriched in differentially expressed proteins. The dotted line corresponds to FDR-corrected P value of 0.05. **e,** *Igf2* is down-regulated at the level of mRNA and protein in 5C embryos, but not placentas at E11.5. Left: RT-qPCR levels of *Igf2* in placenta and embryo at E11.5 (n=6 samples per group). Levels of *Igf2* were normalized against the geometrical mean of *Gapdh*, Sdha and *Pmm1*. Centre: IGF2 protein levels measured by western blot in whole placenta and whole embryo lysates at E11.5 (n=5 per group; Rec – recombinant mouse IGF2 protein; upper and lower arrows indicate the 18 kDa pro-IGF2 and 7.4 kDa mature IGF2, respectively; SOD1 – 19kDa – was used as internal control for loading). Right: quantification of IGF2-total/SOD1 ratios measured by western blot analysis (n=5 samples/group) and shown relative to WT placenta, arbitrarily set to 1. Data are presented as individual values, with averages ± SD in **b, c** and **e** and % indicate ratios 5C/WT; ns – non-significant, ** P<0.01 and *** P<0.001 by two-way ANOVA followed by Šídák’s multiple comparisons tests in **b, c** and **e.**

In a second approach, we engineered an ectopic inducible Tet-off transgene (referred to as *iTg*^Mir483^) (Fig. 4a and Extended Data Fig. 10), the developmental expression of which can be inhibited by the administration of doxycycline (Dox) during pregnancy and beyond (Fig. 4b). Similar to the *Mir483*^5C^ endogenous transgenic model, we found that high levels of *miR-483* expression during intrauterine development (Fig. 4b), caused fetal, but not placental, growth restriction (Fig. 4c) and lethality (Extended Data Fig. 11a). The livers of *iTg*^Mir483^ mutants were disproportionally smaller than in controls (Fig. 4c). The growth restriction of E13.5 fetuses was associated with reduction of IGF2 and IGF1 proteins (Fig 4d). Fetal lethality in *iTg*^Mir483^ was of later onset compared to *Mir483*^5C^ mutants, *i.e*. from E15.5, and was caused by a range of severe defects, in particular malformations of the heart and the great intra-thoracic arteries (Fig. 4e) (see Supplementary Table 2 for full list of developmental abnormalities). Importantly, the lethality and growth phenotypes could be rescued by exposure to doxycycline during pregnancy (Fig. 4f and Extended Data Fig. 11).

**Fig. 4:**
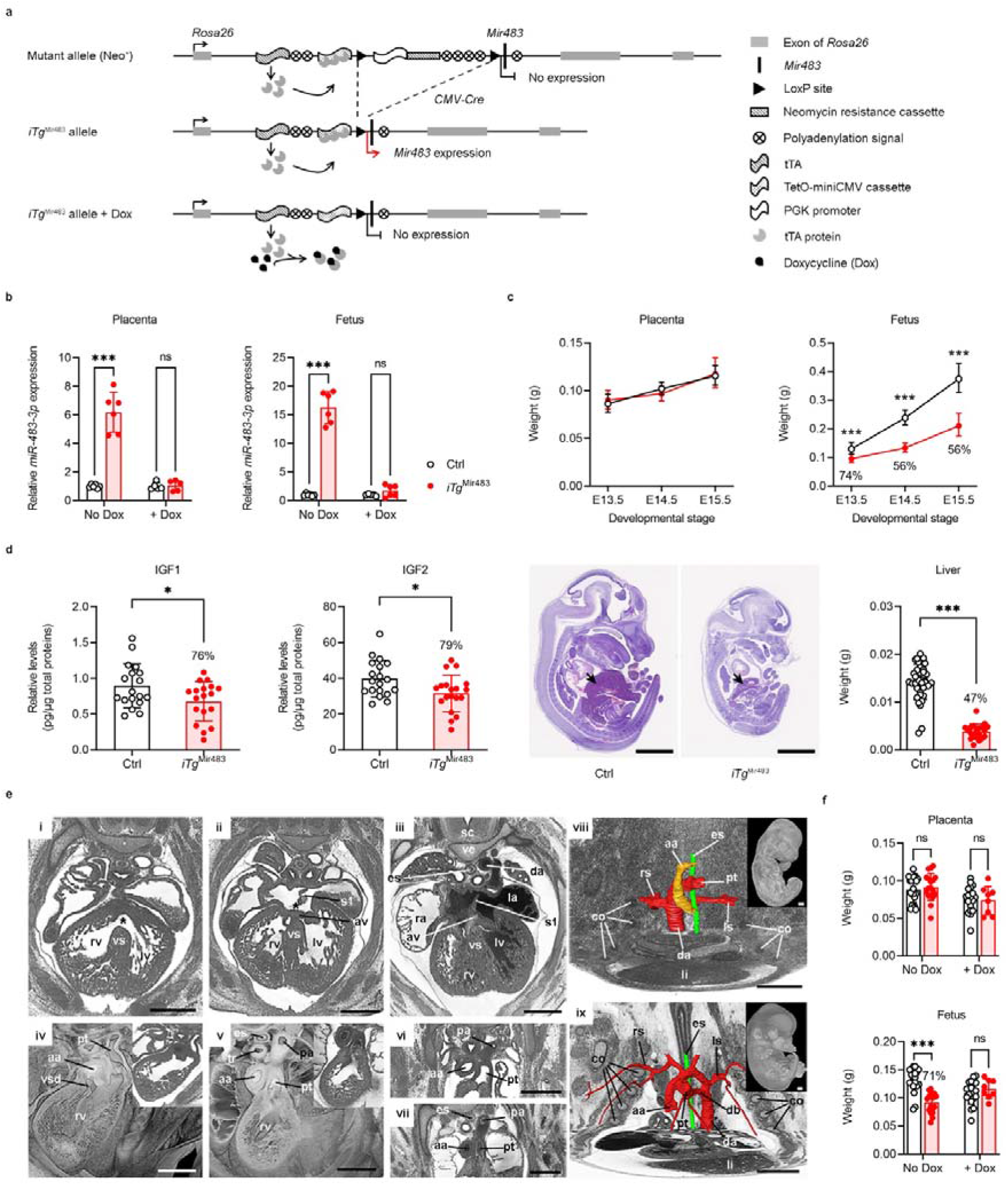
Prenatal developmental phenotyping of a Tet-off inducible over-expressor *Mir483* transgenic model (*iTg*^Mir483^). **a,** Targeting of an inducible *Mir483* transgene by homologous recombination at the *Rosa26* locus. Expression of the *Mir483* transgene is suppressed by the floxed PGK-neomycin-polyA cassettes, which can be deleted *in vivo* in a tissue-specific manner, using Cre-recombinases, and its activity can be controlled by the administration of Doxycycline (Dox) in water, which inhibits the activation of the TetO-miniCMV promoter by the tTA protein (details about the generation of the *iTg*^Mir483^ model are shown in Extended Data Fig. 10). b, Relative levels of *miR-483-3p* expression (normalized against the geometrical mean of *Snord70/snoRNA234* and *Snord68/snoRNA202*) in whole placentae and fetuses at E13.5 without or with Dox exposure (1mg/mL in the drinking water; n=5-6 samples/group). **c,** Without Dox administration, placental growth is normal, while fetal growth is severely compromised, including that of liver, shown at E14.5. The images show representative mid-sagittal sections of E14.5 fetuses (Ctrl – controls; arrows point to the liver, scale bars are 2.5 mm). **d**, IGF1 and IGF2 protein levels measured by ELISA in E13.5 fetuses and normalized against the total protein content measured by a BCA protein assay (n=18-19 samples/group). **e,** Representative high-resolution episcopic microscopy (HREM) scans (n=4 WT and n=6 *iTg*^Mir483^ E14.5 fetuses) identify severe malformations of the heart and the great intra-thoracic arteries, leading to lethality from E15.5 onward: (i-iii) axial HREM-sections showing mutants with septum defects: ventricular septal defect (i, asterisk), atrial septal defect (ii, asterisk) and a WT control (iii); (iv-v) axially-sectioned volume rendered models showing a mutant with a double outlet right ventricle (iv) and a WT control (v); (vi-vii) the central segment of axial HREM sections showing a mutant with coarctation of the aorta (vi) and a WT control (vii); (viii-ix) surface models of intrathoracic arteries and esophagus, illustrating a complex malformation of the great intrathoracic arteries, which combines a right-sided aortic arch, a type B aortic arch interruption and a left sided retroesophageal subclavian artery (viii) and a WT control (ix). The inlay images show entire embryos viewed from the right (note the signs of autolysis in viii). Scale bars are 500 μm; aa, ascending aorta; av, atrioventricular cushion; co, costa; da, descending aorta; db, ductus botalli; es, esophagus; la, left atrium; li, liver; ls, left subclavian artery; lv, ventricle; pa, praeductal aorta; pt, pulmonary trunk; ra, right atrium; rs, right subclavian artery; rv, right ventricle; sc, spinal cord; s1, septum primum; tr, trachea; ve, vertebra; vs, ventricle septum; vsd, ventricle septum defect. **f**, Placental and fetal weights are normal at E13.5 upon Dox administration in the drinking water (1mg/mL) from the beginning of pregnancy (n=8-20 per group). Data are presented as individual values, with averages ± SD in b, c (bottom), d and f, or averages ± 95%CI in **c** (top) and % values indicate ratios *iTg*^Mir483^/Ctrl; ns – non-significant; * P<0.05 and *** P<0.001 by two-way ANOVA followed by Šídák’s multiple comparisons tests in b and f, a mixed effects model in **c** (top), a Mann-Whitney test in **c** (bottom) and unpaired t-tests with Welch’s correction in **d**.

### Growth and metabolic defects in mice with continuous *Mir483* over-expression after birth

In humans, in contrast to mice, both *IGF2* and *MIR483* continue to be expressed in the post-weaning period and throughout adulthood. We exploited the temporal inducible versatility of *iTg*^Mir483^ by extending the postnatal expression of *Mir483* independently of *Igf2* continuously from birth to adulthood, up to 15 weeks of age. We then assessed the consequences for growth, body composition and glucose homeostasis (Fig 5a). Both male and female *iTg*^Mir483^ mice displayed significant post-weaning growth restriction (males 70% of normal; females 86% of normal at 15 weeks of age – Fig. 5b), which resulted from the over-expression of *Mir483* (Fig. 5c). The growth restricted *iTg*^Mir483^ female and male mice had a lower lean mass (Fig. 5d), but showed increased adiposity (Fig. 5d), when compared to controls (Fig. 5e). Fat mass gain in gonadal and subcutaneous depots was particularly extensive at 8 weeks (ranging from 250 to 477% of wild type fat depot weights) compared to 15 weeks (Fig. 5f and Extended Data Fig. 12a), suggesting limited capacity for further adipocyte expansion at the later stage. The increased adiposity was not related to food intake, which was not different in *iTg*^Mir483^ males but in fact, significantly reduced in *iTg*^Mir483^ females (Extended Data Fig. 12b). Adipocytes of *iTg*^Mir483^ were larger compared to controls, as shown *in situ* for the gonadal fat in males (Fig. 5g), and ex *vivo* in both sexes (Fig. 5h), with a notable reduction in the percentage of smaller cells (Fig. 5g and Extended Data Fig. 12c) and the total number of adipocytes per fat pad (Fig. 5g). The increased percentage of larger adipocytes in the gonadal fat was accompanied by down-regulation of *Ttc36* and up-regulation of *Arhgdig* (Fig. 5i), genes recently identified as markers of visceral fat adipocyte hypertrophy in human^32^. However, the increased adipocyte size was not explained by an intrinsic increase in expression of lipid transporters, (Extended Data Fig. 12d), a decreased expression of lipases (which mediate lipolysis, Extended Data Fig. 12e), or an increased expression of enzymes implicated in triglyceride synthesis (Extended Data Fig. 12f). Additionally, protein levels of GDF3, a previously identified target of miR-483-3p^21^, were similar in mature adipocytes of *iTg*^Mir483^ control mice (Denise Fernandez-Twinn *et al*., in preparation). Furthermore, expression of leptin, known to inhibit the expression of adipogenic genes in the white adipose tissue^33^, was up-regulated in adipocytes of *iTg*^Mir483^ mice (Fig. 5j).

**Fig. 5:**
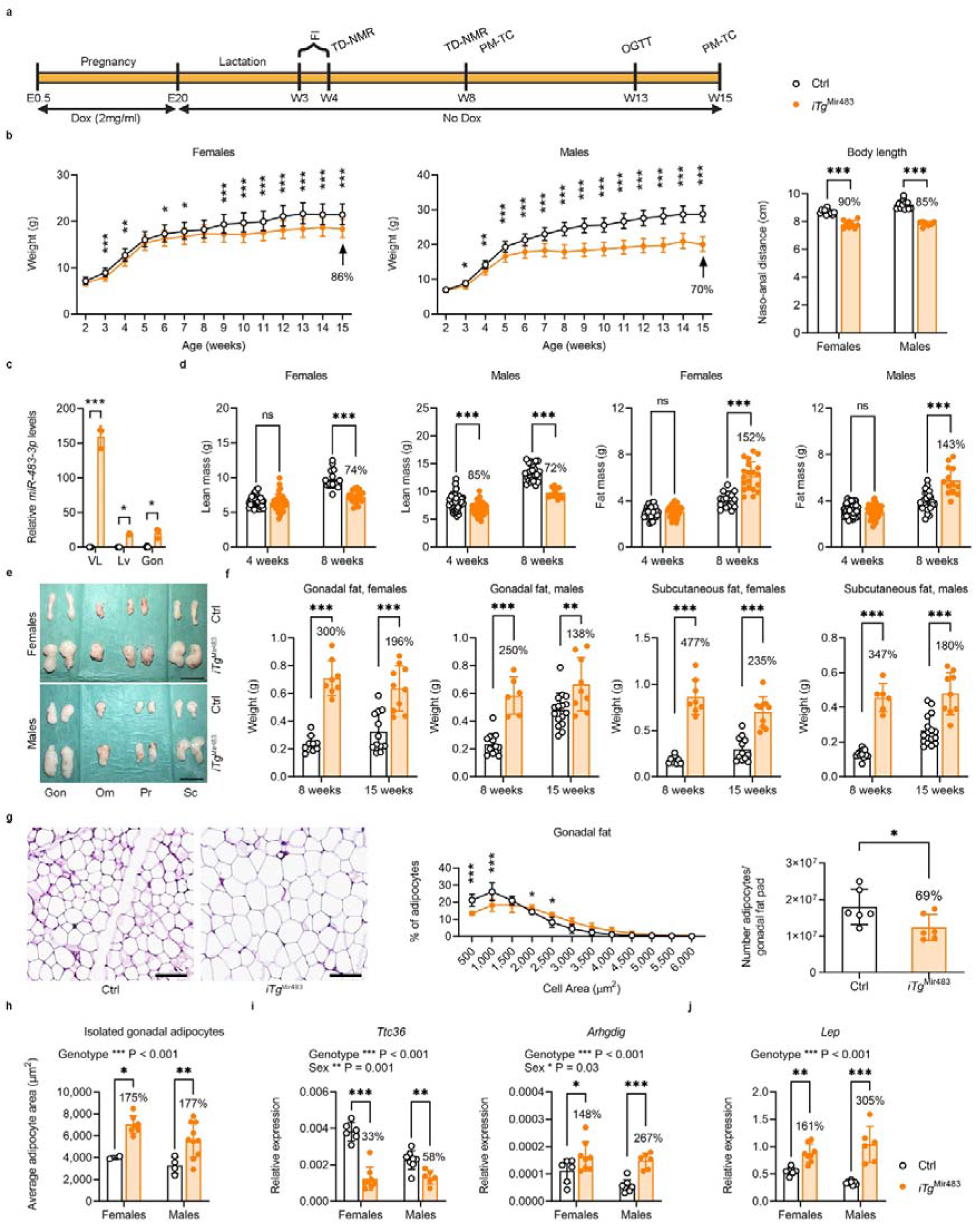
Postnatal *Mir483* overexpression leads to growth retardation, altered body composition, and increased adiposity. **a,** Timeline of the experimental setup: Dox (2mg/ml) was administered in the drinking water throughout pregnancy, after which Dox was withdrawn, allowing expression of *Mir483* from the *iTg*^Mir483^ transgene; food intake (FI) was measured over a period of one week (weeks 3 to 4 – W3-W4); TD-NMR (time-domain nuclear magnetic resonance) was performed at W4 and W8, OGTT (oral glucose tolerance tests) at W13 and post-mortem tissue collection (PM-TC) at W8 and W15. b, *iTg*^Mir483^ mice show postnatal growth restriction in both females (n=10-12) and males (n=9-16) and reduced total body length measured at W15 (n=7-14 mice per sex and genotype). **c**, Relative levels of *miR-483-3p* measured by RT-qPCR in W15 organs (*vastus lateralis* – VL, liver – Lv and gonadal fat – Gon) from *iTg*^mir483^ mutants and littermate controls (Ctrl). Levels of *miR-483-3p* were normalized against the geometrical mean of *Snord70/snoRNA234* and *Snord68/snoRNA202* (n=2-3 samples/group). d, TD-NMR analysis shows reduced lean mass and excessive fat accumulation in *iTg*^Mir483^ mutants compared to littermate controls at W4 and W8 (n=13-51 mice per group). **e,** Individual fat pads are larger in W8 *iTg*^Mir483^ mutants compared to controls: gonadal fat – Gon, omental fat – Om, peri-renal fat – Pr and the sub-inguinal subcutaneous fat – Sc (scale bars are 2 cm). **f**, Gonadal fat pads and sub-inguinal subcutaneous fat pads are significantly heavier in W8 and W15 *iTg*^Mir483^ adults compared to age-matched controls (n=6-16 per group). **g**, Adipocytes are larger in the gonadal fat pads of W15 *iTg*^Mir483^ mutant males compared to age-matched controls (representative H&E stained sections – left, and distribution of adipocyte cell area – middle), with an estimated lower number of mature adipocytes/fat pad – right (n=6 samples per group, scale bars are 100 μm). h, Average cell area measured upon isolation of adipocytes by collagenase digestion from gonadal fat of W8 *iTg*^Mir483^ mutants compared to age-matched controls (n=2-10 samples per group). **i,** Expression patterns of genes that correlate with visceral adipocyte area (*i.e*. reduced *Ttc36* and increased *Arhgdig* mRNA levels) support the adipocyte hypertrophy observed in the gonadal fat of W8 *iTg*^Mir483^ mutants compared to age-matched controls (n=6-8 per group). **j,** Increased expression of Lep gene, encoding leptin in the gonadal fat of W8 *iTg*^Mir483^ mutants compared to age-matched controls (n=6-8 per group). Data are presented as averages ± 95%CI in b (first two graphs on the left), individual values with averages ± SD in **b** (graph on the right), **c, d, f, g** (graph of the right), h, **i** and j, or averages ± SD in g (middle) and values indicate ratios *iTg*^Mir483^/Ctrl and % values indicate ratios *iTG*^Mir483^/Ctrl; ns – non-significant, * P<0.05, ** P<0.01 and *** P<0.001 by a mixed effects model in b (first two graphs on the left), two-way ANOVA followed by Šídák’s multiple comparisons tests in **b** (graph on the right), **c, d, f, g** (graph in the middle), **h, i** and **j** and by an unpaired *t*-test with Welch’s correction in **g** (graph on the right).

We next profiled circulating lipids and observed evidence for a modest dyslipidaemia in both sexes (Fig. 6a). Importantly, we detected drastic reductions in IGF1 levels in both circulation and in organs of *iTg*^Mir483^ mice (Fig. 6b and Extended Data Fig. 13a,b) and increased circulating growth hormone (GH) levels (approximately seven-fold) (Fig. 6c). Given that the increased lipid accumulation in the fat depots and serum dyslipidaemia did not relate to major molecular changes in the adipocytes, we analysed the morphology and function of the liver as a key organ for lipid production. The *iTg*^Mir483^ livers were smaller compared to controls (Fig. 6d), but extensively vacuolated (Fig.6e), suggestive of liver steatosis. The transcription of key genes related to the production of fatty acids, cholesterol and triglycerides were up-regulated in a sex-dependent manner (Fig. 6f-h). Additionally, we found up-regulation of several genes implicated in lipoprotein turnover, as well as key transcriptional regulators of hepatic liver metabolism (Extended Data Fig. 13c). Therefore, our findings suggest that the main site for lipid over-production in the *iTg*^Mir483^ mutants was the liver.

**Fig. 6:**
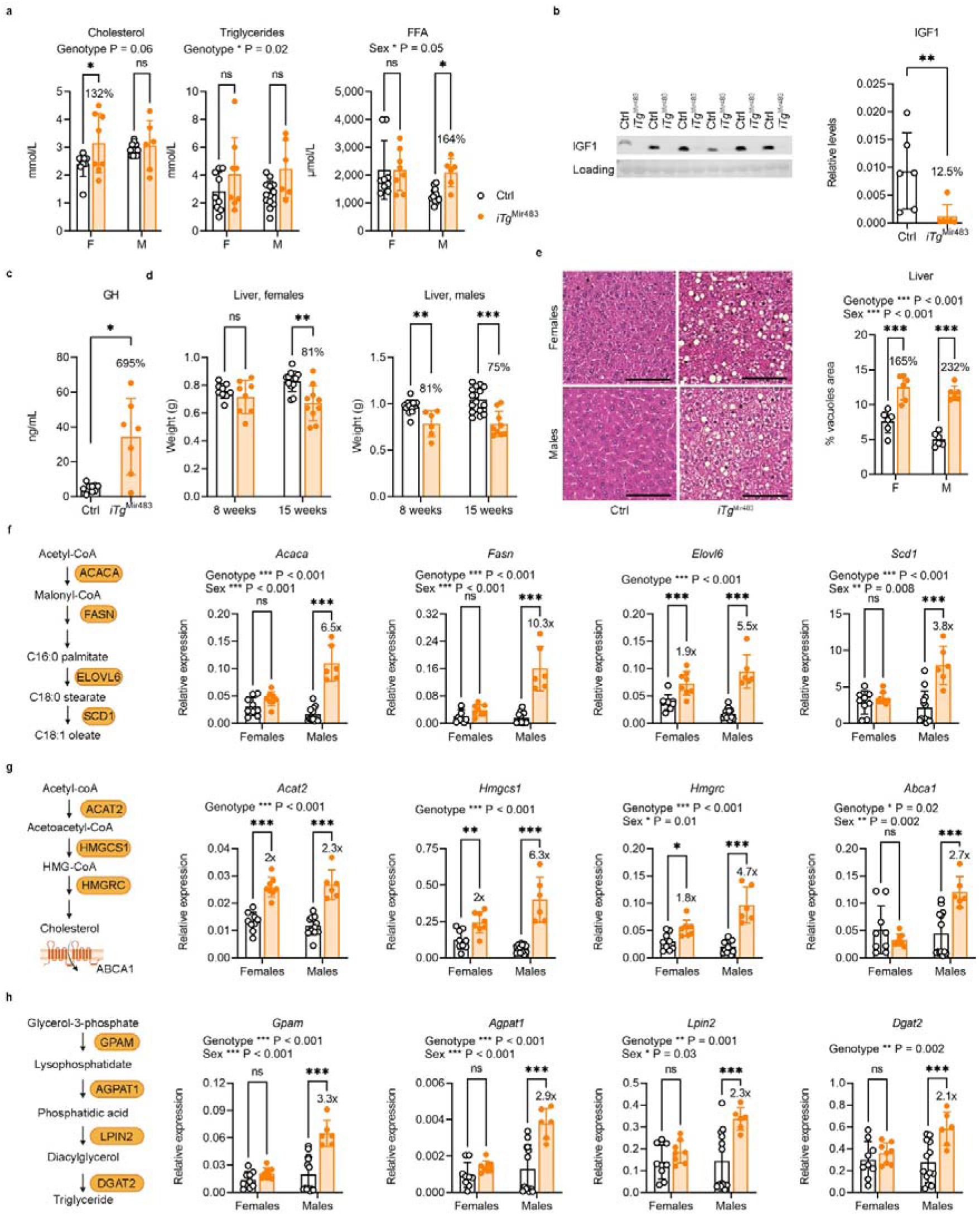
*iTg*^Mir483^ mice show mild dyslipidaemia, altered IGF1 and GH levels, and increased lipid production in the liver. **a,** Serum lipid biochemistry showing moderate dyslipidaemia in W8 *iTG*^mir483^ adults compared to age-matched controls (FFA – free fatty acids; n=6-13 per group). **b**, Circulating plasma IGF1 levels are severely reduced in W15 *iTG*^Mir483^ adults compared to age-matched controls (left – western blotting, right – quantification; n=5 per group). **c**, Circulating plasma GH levels are increased in W15 *iTg*^Mir483^ adults compared to age-matched controls (measurements done by ELISA in n=7-9 per group). **d**, Livers are significantly lighter in W8 and W15 *iTg*^Mir483^ adults compared to age-matched controls (n=6-16 per group), **e,** Livers of W8 *iTG*^Mir483^ adults show significant accumulation of vacuoles, suggestive of liver steatosis (representative H&E stained sections – left, and quantification of % vacuoles area – right; n=6 samples per group, scale bars are 100 μm). f, mRNA levels of genes encoding key enzymes involved in FFA synthesis in the livers of W8 *iTG*^Mir483^ adults compared to age-matched controls (n=6-13 per group). **g**, Relative mRNA levels of genes encoding key enzymes involved in cholesterol synthesis in the livers of W8 *iTG*^Mir483^ adults compared to age-matched controls (n=6-13 per group). **h**, Relative mRNA levels of genes encoding key enzymes involved in triglyceride synthesis in the livers of W8 *iTG*^Mir483^ adults compared to age-matched controls (n=6-13 per group). Data are presented as individual values with averages ± SD and % or fold (x) values indicate ratios *iTG*^Mir483^/Ctrl; ns – non-significant, * P<0.05, ** P<0.01 and *** P<0.001 by two-way ANOVA followed by Šídák’s multiple comparisons tests in **a, d, e, f, g** and **h** or Mann-Whitney tests in **b** and **c**.

The *iTg*^Mir483^ mice showed pronounced multi-organ dysmorphic growth, with disproportional reduction of skeletal muscle and pancreas size, proportionally smaller brain and splenomegaly (Extended Data Fig. 14a). Muscle fibre cell areas were significantly smaller in the *vastus lateralis* of *iTg*^Mir483^ male mice (Extended Data Fig. 14b). Despite these significant changes, glucose homeostasis, assessed by oral glucose tolerance tests (OGTT) in young mice (13 weeks) of both sexes remained normal (Extended Data Fig. 14c). Insulin content of the pancreas was reduced in both sexes (Extended Data Fig. 14d), with a modest tendency for hyperinsulinemia upon fasting (Extended Data Fig. 14e). However, the glucose-induced insulin-secretion (GSIS), assessed during OGTT was impaired, particularly in females (Extended Data Fig. 14e). These data suggest that *iTg*^Mir483^ are insulin-sensitive, despite the increased adiposity, findings which are consistent with the observed normal OGTT profiles. Further evidence in support for an improved insulin sensitivity in *iTg*^Mir483^ mice was provided by the finding of a genotype-dependent increase in pAKT levels in the adipose tissue (Extended Data Fig. 14f).

### Growth defects in *iTg*^Mir483^ mice are rescued by systemic IGF1 infusion

Treating male *iTg*^Mir483^ with a constant infusion of human IGF1 for four weeks (Fig. 7a) rescued whole body growth restriction by the end of treatment (Fig. 7b). However, IGF1-treated *iTg*^Mir483^ did not rescue the excessive fat phenotype (Extended Data Fig. 15a); instead, the gonadal fat pads increased even further in weight (Extended Data Fig. 15b). The brain, liver and *vastus lateralis* also increased in size upon treatment (Extended Data Fig. 15b), with a trend for improved total lean mass (Extended Data Fig. 15a). Further evidence for an *in vivo* link between *miR-483* and IGF1 regulation in the mouse is observed in Igf2^Pat-KO^ and Igf2^+/LacZ^ knockouts^34,35^. In both of these models, which lack both *Igf2* and *Mir483* (Fig. 7c and Extended Data Fig. 15c), *Igf1* mRNA and IGF1 protein levels were increased in fetal liver.

**Fig. 7:**
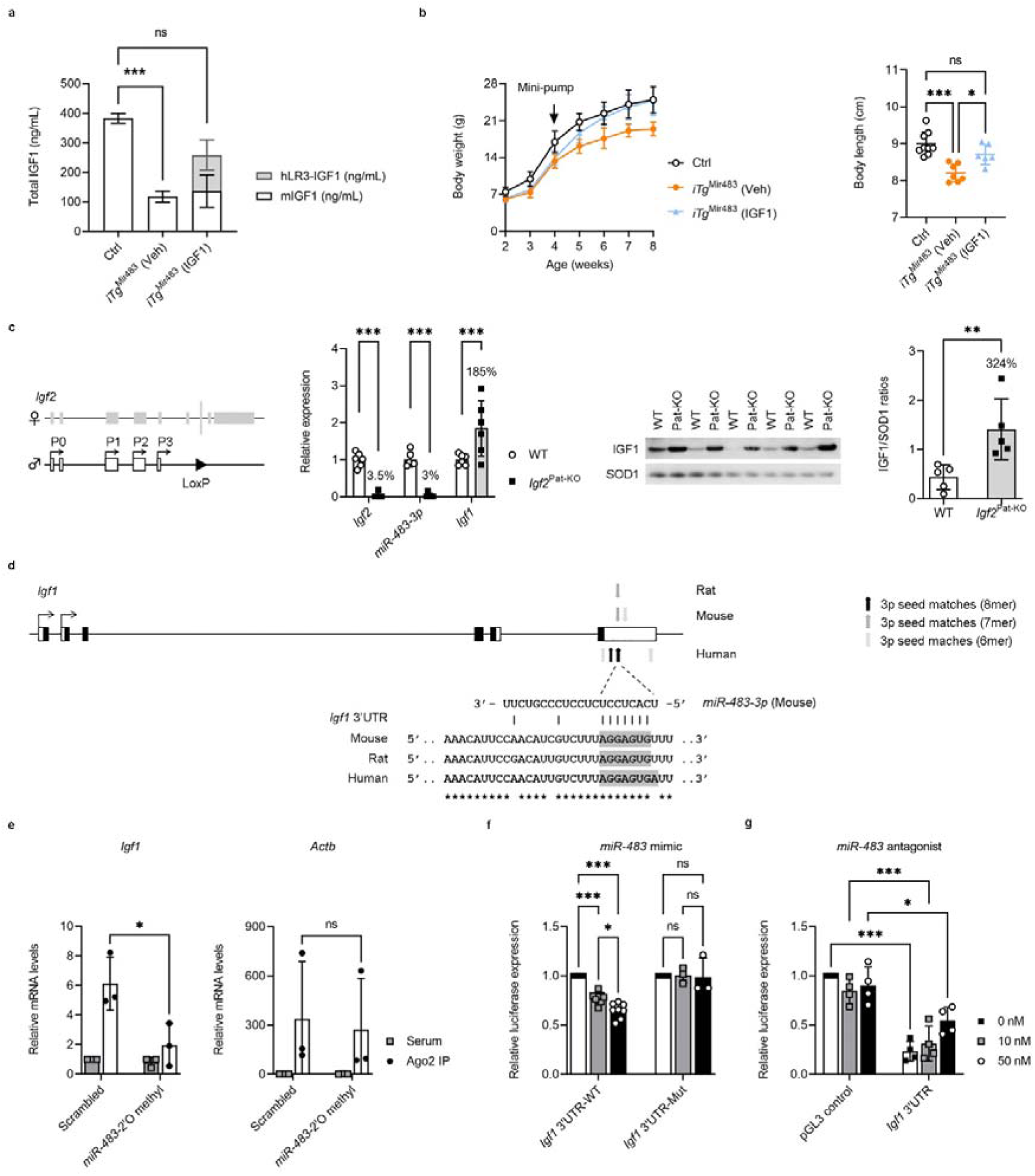
Infusing IGF1 rescues growth phenotypes in *iTg*^Mir483^ mice and IGF1 is a target of *miR-483-3p* both *in vitro* and *in vivo*. **a,** IGF1 infusion via minipump restores the low levels of IGF1 observed in *iTg*^Mir483^ mice; measurements were performed in plasma of W6 male mice, two weeks after the surgery; n=6-7 mice per group. **b**, IGF1 infusion normalizes body weight (left) and body length (right) (n=6-9 male mice per group). **c,** Left: schematic representation of the *Igf2*^Pat-KO^ model in which the coding exons 4-6 of *Igf2* and *Mir483* are deleted using the Cre-loxP technology. Middle: relative RNA levels for *Igf2, miR-483-3p* and *Igf1* in the liver of E18.5 Igf2^Pat-KO^ mutants and WT littermate controls (data was normalized against the geometrical means of *Ppia, Pmm1* and *Hprt* for *Igf2* and *Igf1*, and against the geometrical mean of *Snord70/snoRNA234* and *Snord68/snoRNA202* for *miR-483-3p;* n=6 samples per group). Right: western blot measurement of IGF1 protein in the liver of E18.5 *Igf2*^Pat-KO^ mutants and WT littermate controls, normalized to SOD1 (n=5 per group). **d**, Diagram of the mouse *Igf1* locus showing location of putative *miR-483-3p* seed sequences, and their corresponding locations in the rat and human. The sequence alignment corresponds to the *miR-483-3p* seed-containing sequence in the *Igf1* 3⍰UTR that is conserved across all three species (*miR-483-3p* target sequences are highlighted in grey). **e,** Undifferentiated 3T3-L1 cells were transfected with a *miR-483-* 3p 2⍰-O-Methyl antagonist. Ago2 protein immunoprecipitation was performed, and total RNA was collected. RT-qPCR was used to assay the levels of *Igf1* and *Actb* mRNA present in the immunoprecipitated RISC complex (n=3 independent experiments). **f**, HEK-293 cells (expressing low levels of endogenous *MIR483*) were co-transfected with luciferase reporter constructs containing a portion of the 3’UTR of mouse *Igf1* mRNA spanning the *miR-483-3p* seed target region or mutated seed sequence, together with 0, 10 or 50 nM mouse *miR-483-3p* mimic (n=3-8 samples per group). **g**, HepG2 cells (expressing high endogenous levels of *MIR483*) were co-transfected with luciferase reporter constructs containing a portion of the 3’UTR of mouse *Igf1* mRNA spanning the *miR-483-3p* seed target region, together with 0, 50 or 100 nM *miR-483-3p* 2’-O-Methyl miR antagonist (n=4 samples per group). Data are presented as averages ± SD in a and b (left), individual values with averages ± SD in **b** (right), **c, e, f** and **g** and % values indicate ratios *Igf2*^PatKO^/WT in **c;** ns – nonsignificant, * P<0.05, ** P<0.01 and *** P<0.001 by Kruskal-Wallis test with Dunn’s multiple comparisons tests in a, one-way ANOVA with Tuckey’s multiple comparisons test in **b**, two-way ANOVA followed by Šídák’s multiple comparisons tests in **c** (middle), **e, f**, and **g**, or Mann-Whitney test in **c** (right).

To establish a direct link between *miR-483* and *Igf1*, as a predicted target gene in the mouse (Fig. 7d), we performed Ago2 immunoprecipitation (IP) experiments in undifferentiated 3T3-L1 cells that showed co-localization of IGF1 with the miR processing machinery (Fig. 7e). We then cloned the mouse 3’UTR *Igf1* with either mutated *miR-483* binding sites or wild-type (i.e. non-mutated binding site) controls, into luciferase reporter vectors, and transfected those into low-expressing *MIR483* human HEK-293 cells, in the presence of increasing concentrations of *miR-483* mimic. This led to a decrease in luciferase expression in wild-type 3’UTR transfected cells, which was not observed in 3’UTR mutated transfected cells (Fig. 7f). Conversely, *miR-483* antagonist luciferase-reporter based experiments conducted in human HepG2 MIR483-expressing cells, led to a dose-dependent release of *miR-483* silencing (Fig. 7g).

## Discussion

The Insulin-like growth factor (IGF) signalling system is a major regulator of growth in all vertebrate species, acting through the binding of IGF1 and IGF2 ligands to various receptors to activate pathways involved in cell proliferation, prevention of apoptosis and metabolism^36,37^. *IGF2*, but not *IGF1*, is regulated by genomic imprinting in placental mammals^38^. Here, we provide evidence for a previously unrecognised functional interaction between these two genetic loci to control fetal and postnatal growth. We show that the intronic *Mir483* is co-expressed from the paternal allele with its host gene, *Igf2*, during mouse development, and is a developmental negative modulator of IGF1 levels. This regulatory role was established by several lines of evidence: a) *miR-483* binds to the 3’UTR of the mouse gene and alters IGF1 levels *in vitro;* b) fetuses that lack both *Igf2* and *Mir483* show elevated levels of IGF1 in the liver; c) mice that overexpress *Mir483* exhibit reduced local and systemic IGF1 levels, leading to fetal and postnatal growth restriction; d) systemic infusion of IGF1 in young adult mice that are overexpressing *Mir483* restores the postnatal growth defect. Moreover, work in other species, including human, has revealed that *miR-483* targets *IGF1* in diverse cellular contexts, such as placental cells^39^, NK cells^40^ and myoblasts^41^. In this study, we also provide mechanistic insights into *Mir483* regulation *in vivo:* Mir483 expression is entirely dependent on the *Igf2* transcriptional units and is under the hierarchical control of the gametic imprinting control region ICR1, as shown using the *Igf2*^Δ(P1-P3)^ promoter and H19^δ13/+^ deletion models, respectively.

The uncoupling of *MIR483* from *IGF2* expression in humans seems to have particular importance in human pathology, and in response to environmental cues^42^. In the current study, we aimed to explore the effects of uncoupled expression of *Mir483* from *Igf2* in the postnatal period, using the constitutive inducible TET-off transgene *iTg*^Mir483^. Activation of *Mir483* independent from *Igf2* from birth to adulthood caused postnatal growth stunting, which started around weaning and was more pronounced in males compared to females. A striking adiposity and fatty liver phenotype was associated with the growth impairment of key metabolic organs, suggesting that *miR-483* is a metabolic regulator through yet to be identified target genes, in addition to *Igf1*, which was severely downregulated in most organs and in the circulation. Based on experiments performed on *Igf1* deficient mice models, a phenotype of insulin resistance might be expected^43^ in the *iTg*^Mir483^ over-expressor. However, this was not the case – these mice did not develop glucose intolerance despite increased adiposity and had only low-level dyslipidemia. The absence of a sustained increase in insulin production by the pancreas and the finding of increased pAKT levels in the adipose tissue suggests that these mice have improved adipose tissue insulin sensitivity. The contribution of IGF1 deficiency to some of the *Mir483* phenotypes is clear in terms of whole body and local organ growth, including the effect on lean mass, as IGF1 is a key regulator of muscle mass^44,45^. Infusion of systemic IGF1 in the growth-impaired *iTg*^Mir483^ restored normal growth patterns, thus providing evidence for a causal effect of IGF1 on the growth phenotype. Moreover, reduced levels of IGF1 in *iTg*^Mir483^ may explain the reduced number of mature adipocytes per fat depot, given the well-established role of IGF1 in stimulating pre-adipocyte differentiation^46,47^. How *Mir483* overexpression leads to increased lipid deposition in adipocytes and liver steatosis is currently unknown. Our data suggest that the uptake of lipids in adipocytes may be secondary to the release of excess lipids produced by the liver into the circulation. This is supported by molecular markers of increased lipid production and lipolysis in the liver, but not in the adipose tissue. IGF1 is unlikely to play a role in mature adipocytes, as when pre-adipocytes differentiate they stop expressing IGF1R^46^ (thus, mature adipocytes are not a direct target for IGF1 actions but they secrete IGF1). The differences observed in adipocyte size were independent of the GDF3, previously identified as a direct target of *miR-493-3p*^21^. Identification of additional targets of *miR-483* in the adipose tissue warrants future studies. In human, *miR-483-5p* is positively correlated with the body mass index (BMI), waist circumference, and triglyceride levels and negatively correlated with HDL cholesterol, and is a predictive factor for the development of type II diabetes mellitus and atherosclerosis^48^.

Despite the highlighted differences between mouse and human *Igf2/Mir483* regulation, our work has potential relevance to human imprinting syndromes, cancer and genomic imprinting in general. Children with the intrauterine and postnatal growth retardation Silver Russell syndrome (SRS) that carry DNA hypomethylation epimutations of the ICR1, have decreased levels of IGF2 and increased levels of IGF1 and IGFBP3^49^. Although it remains unknown how *miR-483* levels are regulated developmentally in this syndrome, the increase in IGF1 might be caused by the downregulation of *miR-483* and *IGF2*, as observed in our SRS mouse models, *Igf2*^+/Δ(P1-P3)^ and *Igf2*^+/LacZ^, with elevated IGF1 in fetal liver. Conversely, the overgrowth observed in BWS is associated with increased levels of *IGF2* and *miR-483*^50^, modelled in this study by the H19^Δ13/+^ mice, which also associate elevated *miR-483* levels. Overall, the roles played by *MIR483* in these two human imprinting syndromes remain unclear. Based on our findings we speculate that a key *miR-483* role is the coordination between IGF2 and IGF1 levels, with consequent developmental regulation of growth. We propose that in situations where there are low levels of *IGF2* and of the co-expressed *MIR483*, such as in SRS, IGF1 levels increase to prevent excess fetal growth restriction, thus ensuring fetal viability. Conversely, increased expression of *IGF2* and *MIR483*, as observed in BWS, may lead to decreased IGF1 levels to prevent excessive fetal overgrowth and increased risk for fetal death. Interestingly, during normal human physiology, *IGF2* transcript levels are highest in fetal life compared to postnatal life^38^. The relative drop in *IGF2* transcripts might be accompanied by a reduction in *miR-483* processing in organs such as the liver in the early postnatal period, which may lead to increased levels of IGF1, including circulating IGF1. A systematic study on the regulation of *MIR483* levels in the early postnatal period and adulthood and its correlation with IGF1/IGF2 levels will be required to validate this hypothesis.

*Mir483*, like most imprinted miRs reported to date, is highly expressed in the placenta. Several imprinted miRs play important roles in placenta, including *Mir675*, which also maps to the *Igf2/H19* domain. Mouse studies indicate that processing of *miR-675* from the non-coding *H19* RNA acts to limit placental growth through IGF1R repression^16^. Intriguingly, *Mir483* does not seem to play a role in controlling placental growth, which is consistent with the notion that IGF1 is the main target for regulation, as IGF1 is not expressed in the mouse placenta and IGF1 deficient mice lack a placental growth phenotype^51,52^. It is interesting, however, that the two miRs in – *miR-483* on IGF1 and *miR-675* on IGF1R – the receptor for IGF1 and IGF2. In both cases, they act developmentally as growth suppressors. Although *Mir483* is often referred to as an onco-miR, our work shows that mouse *Mir483* is a growth suppressor (in a developmental context). It is possible that in certain cancer contexts *miR-483* could be used as a therapeutic agent to delay or prevent tumour growth.

Our study has a number of limitations. The overexpression models of *Mir483* are supra-physiological, although the levels of overexpression in the majority of organs in the postnatal period are similar to those normally found *in utero. We* have not addressed the question on why the constitutive knockout of *Mir483* lacks any discernible phenotype. The most likely explanation is the functional redundancy amongst miRs that have key roles in cellular proliferation, as shown in several studies, ranging from mice to worms^53^. Our study did not include environmental manipulations that could stress normal physiology and uncover cellular, molecular and metabolic phenotypes, as reported by several studies. It also remains to be determined the extent to which *miR-483* targets, previously reported in adipocytes^21^, myoblasts^41^, beta cells^25,26^ and hepatocytes^54^, are altered in our inducible model.

In summary, the new loss-of-function and gain-of-function mouse models reported in this study demonstrate that *Mir483* is an imprinted microRNA that acts as a growth suppressor and key regulator of lipid metabolism. The discovery that *Mir483* maintains the balance between two major growth factors, IGF2 and IGF1, supports the concept that this microRNA was evolutionary selected to prevent excessive growth.

## Methods

### Ethics statement

This study was carried out in compliance with the ARRIVE guidelines. The research has been regulated under the Animals (Scientific Procedures) Act 1986 Amendment Regulations 2012 following ethical review and approval by the University of Cambridge Animal Welfare and Ethical Review Body (AWERB). All mouse experiments were approved and performed under PPL No. 70/7594 (study plan 7594/6/15), PPL No. 80/2347 (study plan 2347/2) and PPL No. PC6CEFE59 (study plan IS_AF_001_BF81).

### Nomenclature

Throughout the paper, we are using the current nomenclature for microRNAs, i.e. *Mir483* and *MIR483* when referring to the mouse and human microRNA gene, respectively, *miR-483-3p* and *miR-483-5p* when alluding to the mature miR, and *mir-483* when specifying the stem-loop^14^.

### Generation of the Igf2^Δ(p1-p3)^ mouse model

The *Igf2* gene targeting vector carried a LoxP site and a FRT-flanked neomycin resistance cassette (Neo) inserted 5’ of promoter P1, and a LoxP site inserted 3’ of promoter P3 (Extended Data Fig. 3a). Details of the cloning procedures are available upon request. In brief, we used a 4.0-kb *EcoR*V-*Bci*VI genomic fragment as the 5’ region of homology (5’-ROH), a 5.8-kb *Bci*VI-*Pci*I genomic fragment that includes the P1-P3 promoters as internal ROH, and a 3.9-kb *Pci*I-*Nde*I genomic fragment (intron 3 to exon 6 of *Igf2*) as 3’-ROH. The targeting vector was linearized at a unique *Sca*I site located at the 5’ end of 5’-ROH, and 50 μg linearized vector were electroporated into passage 9, E14 129ola male ES cells, at 250V and 950 μF. Transfected cells were plated onto 10 gelatinized 100-mm dishes preseeded with fibroblast feeder cells. After 24 h in nonselective medium, cells were incubated for 8 days with G418 medium (200 μg/μl) to select for neomycin resistance. Resistant clones were picked at day 9 and expanded into 96-well plates pre-seeded with fibroblast feeder cells. We screened 384 G418-resistant clones by Southern blotting analysis of genomic DNA (gDNA) digested with *Spe*I and hybridized the blots with a unique 635 bp 5’ probe (located external to 5’-ROH and obtained by PCR amplification using primers 5’Pr-F: 5’-CCTGCATAGACGCCTTCCTG-3’ and 5’Pr-R: 5’-GACCCTAACTCTCCCAAGTCCC-3’) (Extended Data Fig. 3b). Two correctly targeted clones at the 5’ end were then verified by Southern blot (*EcoR*I digested DNA) using a 733 bp 3’ probe (located external to 3’-ROH and obtained by PCR amplification using primers 3’Pr-F: 5’-GCCCAAGTAACCTGACCCCT-3’ and 3’Pr-R: 5’-CGAGCACCTTCCTAACACCTG-3’) and an additional check for multiple integrations elsewhere in the genome using a 628 bp internal probe (located in *Igf2* promoter P1 and obtained by PCR amplification with primers Int-F: 5’-CCACCACATTTAGACAGCATT-3’ and Int-R: 5’-ACCGTAGGAGAAGTGACGAG-3’). Two clones with a single integration site and correctly targeted 5’ and 3’ LoxP sites were thus identified (Extended Data Fig. 3c), with the loxP sequences further verified by Sanger sequencing. The Neo cassette was excised by transiently transfecting the two ES cell clones with FLPe recombinase, followed by two rounds of subcloning. Four correctly excised clones that carry a single FRT site and lack the Neo cassette were identified by PCR screening (288 ES subclones) using primers F: 5’-ATGTCTCCAATCCTTGAACACTG-3’ and R1: 5’-GCAGTGGGAGAAATCAGAACC-3’ (Extended Data Fig. 3d). Two independent ES clones were then microinjected into C57BL/6J blastocysts and transferred into (C57BL/6J X CBA/Ca) F1 pseudopregnant females to generate chimeric mice. Four chimeras were born, two males and two females and germline transmission was achieved only through the females. Germline transmitting mice were backcrossed into the C57BL/6J genetic background for more than 10 generations before being used as experimental animals, and genotyping was performed by PCR using primers listed in Supplementary Table 3 (Extended Data Fig. 3e). Efficient deletion of the floxed *Igf2* P1-P3 region upon paternal transmission of the targeted allele was verified by PCR (Extended Data Fig. 3f) and by Northern blotting (Extended Data Fig. 3g).

### Generation of a *Mir483* specific knockout mouse

Targeted JM8.F6 ES cells (from the C57BL/6N mouse strain) (International Knockout Mouse Consortium Project Design ID: 49935) were provided by Dr. Haydn Prosser^55^. Confirmation of targeting was performed by Southern blotting (Extended Data Fig. 6). The 5’ probe (587 bp, generated by PCR amplification using primers 5’arm_F2: 5’-GGCTTACTGTGGGTCATCGT-3’ and 5’arm_R2: 5’-CTGGACACTGGACCTGGTTT-3’) was hybridised to *Eco*RV-digested DNA to give expected bands of 12,984 bp (targeted locus, Mut) and 18,532 bp (wild-type locus, WT). The 3’ probe (685 bp, generated by PCR amplification using primers 3’arm_F: 5’-ATGTGTGACCAGGCTGCTAGTTC-3’ 3’arm_R: 5’-GTGTTGATGGCTCTAGCTGGTGT-3’) was hybridized to *Xba*I-digested DNA to give expected bands of 3,562 bp (Mut) and 10 447 bp (WT). A single insertion of the targeted vector was confirmed using the Puro probe (537 bp, generated by PCR amplification using primers Puro_F: 5’-GGTCACCGAGCTGCAAGAAC-3’ and Puro_R: 5’-AGTTGCGTGGTGGTGGTTTT-3’) hybridized to *Eco*RV-digested DNA, which gives the band of 12,984 bp (Mut), or to *Xba*I-digested DNA, which gives the expected band of 9,104 bp (Mut).

ES cells were cultured using standard conditions as recommended by the International Knockout Mouse Consortium. Selection marker was excised by transient in vitro transfection with pCAG-Cre (a gift from Dr. Connie Cepko – Addgene plasmid 13775^56^) encoding a Cre recombinase and ES cells were then genotyped by PCR using primers F1: 5’-TACCTGCCTGTGAACTGCTCTG-3’, R1: 5’-ATCTGGTGCCTCCTGTCTGGTA-3’ and R2: 5’-CTCTGAGCCCAGAAAGCGAAG-3’ (expected product sizes: PGKPuroΔTK allele – 560 bp, WT allele – 440 bp, *Mir-483*^KO^ allele – 457 bp). *Mir-483*^KO^ ES cells from two independent clones (F4 and E12) were injected into C57BL/6J-Tyrc-2J (albino) blastocysts. Chimeric mice (identified by their coat colour) were then mated with albino B6 mice and germline transmission was validated by the appearance of black offspring. Subsequently, the colony was maintained by crossing with C57BL/6J wild-types. Mice were genotyped by PCR using primers F1 and R1 (Extended Data Fig. 6).

### Generation of *Mir483*^5C^ mice with five copies of *Mir483* at the endogenous locus

To generate the plasmid for recombination-mediated cassette exchange (RCME), we followed the following steps (Extended Data Fig. 8). (1) First, one copy of *Mir483* with 150 bp of flanking sequence was amplified using primers 102_F: 5’-ccccccctcgaggtcgacggtatcgatTCTTCACTTCTGCCTACCTGCCTG-3’ and 107_R: 5’-CGGGCTGCAGGAATTCagtggtttggaaaacagggaggag-3’. The PCR product thus obtained was cloned using the In-Fusion HD Cloning Plus kit (Takara Bio – 638909) into the pMA_F3NeoLoxPPolyFRT vector provided by Dr. H. Prosser^54^, linearized with *Cla*I and *Eco*RI. The resulting one-copy plasmid was digested with *Eco*RI and *Bam*HI and two further copies of *Mir483* (generated by PCR using primers 103: 5’-AAGCTTAGTGGTTTGGAAAACAGGGAGGAG-3’ plus 108: 5’-CCAAACCACTGAATTTCTTCACTTCTGCCTACCTGCCTG-3’ and 104: 5’-CAAACCACTAAGCTTTCTTCACTTCTGCCTACCTGCCTG-3’ plus 109: 5’-TAGCCCGGGCGGATCCAGTGGTTTGGAAAACAGGGAGGAG-3’) were cloned in, using the In-Fusion HD Cloning Plus kit. The resulting three-copy construct was further digested with *Bam*HI and two additional copies of *Mir483* (generated by PCR using primers 112: 5’-CCAAACCACTGGATCCTCTTCACTTCTGCCTACCTGCCTG-3’ plus 105: 5’-AGGCAGAAGTGAAGAAGTGGTTTGGAAAACAGGGAGGAG-3’ and 106: 5’-GTTTTCCAAACCACTTCTTCACTTCTGCCTACCTGCCTG-3’ plus 109: 5’-TAGCCCGGGCGGATCCAGTGGTTTGGAAAACAGGGAGGAG-3’) were additionally cloned-in to produce the five-copy plasmid. (2) To perform the RCME and screening, the five-copy plasmid obtained above was mixed with the pPGKFLPobpA plasmid (a kind gift from Dr. Philippe Soriano – Addgene plasmid 13793^57^), encoding FLPo recombinase and the mix was co-electroporated into ES cell clones which had confirmed replacement of *Mir483* with PGKPuroΔtk allele (Extended Data Fig. 8). Single-cell-derived colonies were picked and screened by PCR using primers F2: 5’-GTGCCACTCCCACTGTCCTT-3’ and R1: 5’-ATCTGGTGCCTCCTGTCTGGTA-3’, generation of a 2,430 bp product indicating that RCME had occurred and five copies of *Mir483* were present. The identity of the PCR product was further confirmed by digestion with *Bam*HI, *Hind*III or *Cla*I (Extended Data Fig. 8). (3) The intermediate Neo cassette was deleted in vitro by transfection with pCAG-Cre (a gift from Dr. Connie Cepko – Addgene plasmid 13775^56^) encoding a Cre-recombinase. ES cells were genotyped using primers 142: 5’-CACGCTTCAGTTTGTCTGTTCG-3’, 144: 5’-CGTGCTACTTCCATTTGTCACG-3’ and 145: 5’-CTGGAGTGGTTTGGAAAACAGG-3’, which generate products of 1,014 bp (Neo^+^ allele) and 925 bp (WT allele), and also using primers 142, 145 and 143: 5’-AAGAATCGATACCGTCGACCTC-3’, which generate products of 740 bp (Neo^−^ allele) and 925 bp (WT allele). This latter set of three primers was also used for genotyping *Mir-483*^5C^ mice (Extended Data Fig. 8).

### Generation of *iTg*^Mir483^ mice with an additional copy of *Mir483* inserted at the *Rosa26* locus

To generate the targeting vector containing one-copy of *Mir483* with Tet regulation (*Mir483*^1c^), *Mir483* was amplified from the one-copy plasmid made for RCME, using primers 135: 5’-TGCAGCCCAAGCTAGCCCCTCGAGGTCGACGGTATCGAT-3’ and 136: 5’-GCGGGGGCCCCTCGAGCTCCACCGCGGTGGCGGCCGCTC-3’ and the resulting 485 bp product was cloned between *Nhe*I and *Xho*I restriction sites of the pTET-BigT plasmid (a kind gift from Dr. Andrew P. McMahon^58^). To enable ES cell targeting at the *Rosa26* locus, the pTET-BigT-*Mir483*^1c^ construct was digested with *Pac*I and *Asc*I and subcloned into pROSA26PAS (a kind gift from Dr. Andrew P. McMahon^58^). pROSA26PAS-*Mir483*^1c^ was linearised with *Afe*I and electroporated into BayGenomics E14Tg2A.4 ES cells and JM8.F6 ES cells.

ES cell clones were screened by PCR. Homologous recombination of the 5’ arm was assayed using primers 146: 5’-CGCCTAAAGAAGAGGCTGTG-3’ and 148: 5’-GAAAGACCGCGAAGAGTTTG-3’ (expected product size of 1,316 bp). Homologous recombination of the 3’ arm was assayed using primers 152: 5’-GGGAGGATTGGGAAGACAAT-3’ and 153: 5’-CGAAGACCTGTTGCTGCTCA-3’ (expected product size of 4,779 bp) (Extended Data Fig. 10).

Targeted ES clones and the subsequent transgenic mice were also genotyped using primers 147, 148, 156, which generate 326 bp product from *Tg* (Neo^+^) allele and 436 bp from the WT allele (Supplementary Table 3). In vivo Cre-mediated deletion of the Neo cassette was determined using primers 151 and 143, which generate a 266 bp product from the deleted allele (*iTg*^Mir483^), or a 2,931 bp product if deletion does not occur (Supplementary Table 3). Presence of Cre-recombinase was confirmed using primers Cre-F and Cre-R and Ctrl-F and Ctrl-R, which generate a 390 bp product from the *Cre* transgene and a control 254 bp product (Extended Data Fig. 10).

### Additional mouse strains and mouse husbandry

*H19^Δ13^* mutant mice^30^, *Igf2*^LacZ^ mice^35^ and *CMV-Cre* mice^59^ were generated previously and were obtained from the Babraham Institute, Cambridge. The Igf2^fl/fl^ mice were generated in our laboratory, as previously described^34^. The *CMV-Cre* recombinase is expressed soon after fertilization and allows ubiquitous deletion of floxed alleles in all tissues, including the germline^59^. C57BL/6J mice used as wild-type (WT) controls were purchased from Charles River (Strain Code: 632).

All mouse work was performed unto a C57BL/6J genetic background. Mice were maintained and mated under pathogen-free conditions at the University of Cambridge Phenomics Unit (West Forvie). They were fed a standard chow diet with 9% of kcal from fat (SDS, Essex, UK), and housed with a 12-h light/dark cycle in a temperature-controlled room (22°C). Food and water were available ad libitum, except for periods of fasting when food was withdrawn. For timed mating, the day of detection of a vaginal plug was noted as embryonic day 0.5 (E0.5) and the day of birth was noted as post-natal day 0 (P0). Mice were weaned at 3 weeks of age and ear notches were used for visual identification and genotyping, which was performed using standard PCR, with primers listed in Supplementary Table 3, followed by separation of PCR amplicons by agarose gel electrophoresis.

### Southern blotting

Southern blotting analyses of genomic DNA extracted from ES clones was performed as previously described^34^. Briefly, genomic DNA were digested with appropriate restriction enzymes, then DNA fragments were separated by electrophoresis on 0.8% agarose gels in 1×TBE buffer, alkaline blotted onto Hybond N+ membranes (Amersham), and UV cross-linked (Stratalinker, Stratagene). Probes were obtained by PCR as described above and radiolabelled (α-32P-CTP). After hybridisation and washing, the membranes were exposed overnight to MS film (Kodak).

### Northern blotting

Northern blotting analysis of Igf2 expression in E18.5 placenta and liver samples was performed as previously described^34^. Briefly, total RNA (10 μg) was separated in low-percentage formaldehyde-treated agarose gels, blotted onto Nytran-plus membrane (Schleicher and Schuell), and UV crosslinked (Stratalinker, Stratagene). The RNA blots were hybridized with radiolabelled (α-32P-UTP) *Igf2* and *Gapdh* (internal control) cDNA probes. After hybridization and washing, transcript levels were quantified by PhosphorImager analysis (Molecular analyst software, Biorad).

### Sequence alignments

Sequences of eutherian mammals and marsupials corresponding to *Mir483*, its promoter and target sequences at the *Igf2* 5’UTR and *Igf1* 3’UTR were retrieved from NCBI (National Center for Biotechnology Information). These sequences were then aligned using Clustal Omega (https://www.ebi.ac.uk/Tools/msa/clustalo/), using ClustalW as output format.

### RNA extraction and RT-qPCR

Total RNA was extracted from tissues or cells using RNeasy Plus Mini Kits (Qiagen – 74134). Small RNAs were extracted using mirVana kits (ThermoFisher Scientific - AM1560) or miRNeasy Mini kits (Qiagen – 217004). Total RNA was treated with an RNase-Free DNase Set (ThermoFisher Scientific – EN0521). RNA concentrations were measured by NanoDrop (Thermo Scientific) and quality was assessed in 1.2% agarose gels, or using the RNA 6000 Pico or Nano Kits (Agilent – 5067-1513 and 5067-1511) and an Agilent 2100 Bioanalyzer. Total RNA (200 ng) was reverse transcribed into cDNA using the RevertAid RT Reverse Transcription Kit (ThermoFisher Scientific – K1622). For microRNA, reverse transcription was performed using the TaqMan MicroRNA Reverse Transcription Kit (ThermoFisher Scientific – 4366596) from 4.8ng total small RNA.

RT-qPCR was performed with the SYBR Green JumpStart Taq Ready Mix (Sigma – S4438) and custom-made primers (Supplementary Table 4), or with TaqPath ProAmp Master Mix (ThermoFisher Scientific – A30866) and TaqMan probes (Supplementary Table 4) in an ABI Prism 7900 system or QuantStudio6 Real-time PCR machine (Applied Biosystems). Gene expression normalisation was performed against combination of the housekeeping genes: *Ppia* (peptidylpropyl isomerase A or cyclophilin-A), *Gapdh* (glyceraldehyde 3-phosphate dehydrogenase), *Pmm1* (phosphomannomutase 1), *Hprt* (hypoxanthine phosphoribosyltransferase), *Actb* (actin beta), *Tbp* (TATA box binding protein) and *Sdha* (succinate dehydrogenase complex flavoprotein subunit A), as appropriate. For small RNAs, expression levels were normalized against *Snord70/snoRNA234* and/or *Snord68/snoRNA202*, used as internal controls. Relative levels of expression were calculated using the 2^−ΔΔCt^ method^60^.

### Food intake

Food intake was measured in the *iTg*^Mir483^ model between W3 (week 3) and W4. Briefly, two mice of same sex and genotype were placed in a new cage and the food pellets were weighted at 4 pm at the start and the end of the seven-day interval. The average food consumed (g/day/mouse) was calculated by dividing the food consumed per cage in a week by 14.

### Body composition

For the *Mir483*^KO^ model, body composition was analysed by DEXA scanning (Lunar PIXImus densitometer, General Electric) immediately after killing by cervical dislocation between 8 am and 10 am. For fat mass and lean mass, values were expressed as a proportion of total body weight, and the bone mass density (BMD) was calculated related to the body length (naso-anal distance) and presented as g/cm^2^. For the *iTg*^Mir483^ model, body composition analysis was performed at W4 and W8 on live and conscious mice, using time-domain nuclear magnetic resonance spectroscopy (TD-NMR) with the Minispec Live Mice Analyser (Bruker Minispec Live Mice Analyser LF50) that measures total body fat mass and lean mass^61^. For *iTg*^Mir483^ mice that underwent surgery (minipump insertion), the TD-NMR at W8 was performed post-mortem, immediately after the removal of the empty minipump.

### Histology and stereology analyses

Immediately after dissection, tissues (gonadal fat, *vastus lateralis* skeletal muscle and liver) were fixed in 10% buffered formalin for 48 hours, then were dehydrated and embedded in paraffin. Paraffin blocks were cut at 5 μm thickness, sections were then deparaffinised, rehydrated, stained (using a standard haematoxylin-eosin staining protocol) and mounted with coverslips. For all stereological analyses, in order to obtain accurate morphometric estimations, at least two sections/block, spaced at 200 μm, were used. The stained slides were imaged using the Zeiss Axioscan Z1 Slidescanner (Carl Zeiss). Whole-slide scans of stained sections were analysed using HALO or HALO AI (Indica Labs) to measure the area of individual adipocytes, skeletal muscle fibres or the percentage occupied by vacuoles in the liver.

### Glucose tolerance tests

Glucose tolerance tests were performed as previously described^62^. Briefly, for the *Mir483*^KO^ model, glucose was administered at 1 mg/g body weight by intra-peritoneal injection (ipGTT) after 16 h fasting (5 pm to 9 am the following day) at the age of 12 weeks. For the *iTg*^Mir483^ model, glucose was administered at 2 mg/g body weight by oral gavage (OGTT) after 6 h fasting (8 am – 2 pm) at the age of 13 weeks. The areas under the curve (AUCs) following ipGTTs and OGTTs were calculated by the trapezoidal rule, after normalization to basal glucose levels.

### Tandem Mass Tag (TMT) analysis

This analysis was performed using the TMTsixplex Isobaric Mass Tagging Kit (ThermoFisher Scientific – 90064) that allows multiplexing up to six samples. Three WT and three 5C embryos at E10.5 were first suspended in RIPA buffer (ThermoFisher Scientific – 89901) and dissociated using Dounce homogenizers. Protein concentrations were determined using a BCA assay (ThermoFisher Scientific – 23221). Once quantified, 100 μg protein per condition were transferred, to a final volume of 100 μL with 100 mM TEAB (triethylammonium bicarbonate buffer). Then, 5 μL of the 200 mM TCEP (tris(2-carboxyethyl)phosphine) was added to every sample, which were then incubated at 55°C for 1 hour. The samples were incubated for additional 30 minutes and protected from light once 5 μL of the 375 mM iodoacetamide were mixed in. The proteins were then precipitated with excess pre-chilled (−20°C) acetone and incubated overnight at −20°C. Samples were centrifuged (8000 × *g* for 10 minutes at 4°C), the supernatant removed and the pellet air-dried. Subsequently, the pellet was resuspended with 100 μL of 100 mM TEAB. In addition, 2.5 μL of 1 μg/μLtrypsin solution (*i.e*. 2.5 μg) were added to each protein sample, followed by overnight digestion at 37°C. 41 μL of anhydrous acetonitrile were added to each of the mass tag (with reporter ions from m/z = 126.1 to 131.1) and each merged with one protein sample, in no particular order. The reaction was incubated for 1 hour at room temperature and quenched with 8 μL of 5% hydroxylamine for further 15 minutes. Finally, the samples were pooled at equal concentrations and stored at −80°C until mass spectrometry analysis.

The multiplexed TMT samples were submitted to the CIMR/IMS Proteomics Facility (CIPF) for fractioning and mass spectrometry. Data was collected using a Q Exactive Orbitrap Mass Spectrometer with EASY-spray source and Dionex UltiMate 3000 RSLC System (ThermoFisher Scientific). The software search engine MASCOT (Matrix Science) was used to identify proteins.

### Protein extraction and western blotting

Protein was extracted in lysis buffer (50 mmol/l HEPES [pH 8], 150 mmol/l NaCl, 1% (wt/vol.) Triton X-100, 1 mmol/l Na_3_VO_4_, 30 mmol/l NaFl, 10 mmol/l Na_4_P_2_O_7_, 10 mmol/l EDTA (all Sigma–Aldrich) with a cocktail of protease inhibitors [set III, Calbiochem, Merck Life Science UK Ltd.]. Total protein concentration of lysates was determined using a bicinchoninic acid kit (Merck Life Science UK Ltd, Gillingham, UK) and samples diluted in Laemmli buffer. Total protein from tissue extracts or plasma were prepared in Laemmli or RIPA buffers (both from Merck Life Science UK Ltd, UK), separated by polyacrylamide gel electrophoresis and then transferred to either nitrocellulose or a polyvinylidene difluoride (PVDF) (Merck Life Science UK Ltd, UK). Membranes were then processed for Western blotting using the antibodies listed in Supplementary Table 5. Protein bands were visualised using a chemiluminescence substrate (Immobilon Forte, Merck Life Science UK Ltd.) on the ChemiDoc Imaging system (Bio-Rad, Hemel Hempstead, UK). Protein abundance was quantified by band densitometry, using Image Lab 6.1 software (Bio-Rad), and normalised to levels of SOD1 or the total protein transferred, assessed by Coomasie-250 staining.

### IGF1, IGF2 and GH measurements by ELISA

All measurements were performed at CBAL (Core Biochemical Assay Laboratory, Addenbrooke’s hospital). IGF1 was measured using the mouse/rat IGF-1 Quantikine ELISA kit (Biotechne – MG100), which employs the quantitative sandwich enzyme immunoassay technique. Briefly, a monoclonal antibody specific for mouse/rat IGF-1 was pre-coated onto a microplate. Standards, controls and samples were pipetted, in duplicate, into the wells and any IGF-1 present became bound by the immobilized antibody. After washing, an enzyme-linked polyclonal antibody specific for mouse/rat IGF-1 was added to the wells. Following a wash to remove any unbound antibody-enzyme reagent, a substrate solution was added to the wells and colour developed in proportion to the amount of IGF-1 bound in the initial step. The colour development was stopped and the intensity of the colour was measured on the Perkin Elmer Victor3 plate reader. Samples were assayed in duplicate on a 1:500 dilution.

GH was measured using a mouse/rat growth hormone ELISA kit (Millipore – EZRMGH-45K), based on quantitative sandwich enzyme immunoassay technique. Briefly, a polyclonal antibody specific for mouse/rat growth hormone was pre-coated onto a microplate. Standards, controls and samples were pipetted, in duplicate, into the wells and any growth hormone present became bound by the immobilized antibody. After washing, a biotinylated polyclonal antibody specific for mouse/rat growth hormone was added to the wells. After further washing, a streptavidin-horseradish peroxidase conjugate was added to the wells. Following a wash to remove any unbound conjugate, a substrate solution was added to the wells and colour developed in proportion to the amount of growth hormone bound in the initial step. The colour development was stopped and the intensity of the colour was measured at 450nm on the Perkin Elmer Victor3 plate reader. Samples were assayed in duplicate, using 10 μl undiluted plasma.

IGF2 measurements in E13.5 whole-embryo lysates were performed with the Mouse IGF-II DuoSet ELISA kit (R&D Systems – DY792), using an assay adapted for the MesoScale Discovery electrochemiluminescence immunoassay platform (MSD), as previously described^63^.

### Plasma insulin and total-pancreas insulin measurements

Blood samples for plasma insulin measurements were collected from the tail vein in heparinised capillary tubes during the OGTT experiments at 0 and 20 min, from W13 *iTg*^Mir483^ mouse model. The tubes were kept on ice and spun at 4,000 RPM (rotations per minute) for 5 min. Plasma samples were flash-frozen in liquid N2 and stored at −80°C until analysis. For total-pancreas insulin measurements, the entire pancreases collected from W8 *iTg*^Mir-483^ mouse model were collected into cold acid-ethanol (0.18M hydrochloric acid in 70% (vol/vol) ethanol and flash frozen in liquid N2, then pulverised and sonicated, stored at 4°C overnight before storage at −70°C until analysed. Insulin levels in plasma were measured using ELISA kits (Meso Scale Discovery Mouse/Rat Insulin Assay Kit) at CBAL (Core Biochemical Assay Laboratory, Addenbrooke’s hospital). Insulin levels in acid–ethanol supernatants were measured using ELISA kits (Mercodia – 10-1247-01). Total pancreas insulin content (pmol/L) was normalised to the total pancreas wet weight (g), measured at collection.

### Blood biochemistry

Serum cholesterol, triglycerides and free fatty acids concentrations were measured using enzymatic assay kits, as previously described^62^. Briefly, total cholesterol was measured using an enzymatic assay kit (Siemens Healthcare – DF27) that combines activities of cholesterol esterase and cholesterol oxidase. Triglycerides were measured using an enzymatic assay kit (Siemens Healthcare – DF69A) that combines activities of lipoprotein lipase, glycerol kinase and glycerol-3-phosphate oxidase. The assays for total cholesterol and triglycerides were automated on the Siemens Dimension EXL analyser. Free (non-esterified) fatty acids were measured using Roche’s Free Fatty Acid Kit (half-micro test) (Sigma Aldrich – 11383175001), which is based on the enzymatic conversion of free fatty acids to acyl CoA by acyl-Co A synthetase.

### High-resolution episcopic microscopy (HREM)

E14.5 fetuses were fixed, dehydrated, infiltrated and embedded as previously described^64^. Embedded fetuses were analysed by HREM, using 3-μm sections, green fluorescent protein filters and a Hamamatsu Orca HR CCD camera to obtain the high-resolution images, as previously shown^65^, and datasets were analysed with the Amira 5.4 software (Visage Imaging). For illustration, volume rendering was combined with arbitrary section plane erosion in Amira, to obtain the fetus models shown in Figure 4. Individual aortae, pulmonary arteries and esophagus were also digitally segmented from image stacks in Amira, and used to generate surface-rendered pseudo-coloured 3D organ models, which were superimposed in the appropriate location upon semi-transparent volume rendering of the fetus.

### Ago2 immunoprecipitation

Undifferentiated 3T3-L1 cells were transfected with 100 nM of mouse *miR-483-3p* 2⍰-O-Methyl antagonist, or a scrambled 2⍰-O-Methyl RNA sequence (both custom-made, Sigma). The cells were harvested in PBS, and then fixed in the presence of 1% formaldehyde for 1⍰hour, to establish protein-RNA reversible crosslinks. Cells were lysed and sonicated, then the protein lysate was immunoprecipitated using a specific Ago2 mouse monoclonal antibody (Abcam – ab186733) and bound by A/G agarose beads (Santa Cruz Biotechnology – SC-2003). Beads bound to the antibody were resuspended in water after precipitation and heated at 75°C for 45⍰min to disrupt the RNA-protein interactions. The RNA was then extracted and purified using Trizol. *Igf1* and *Actb* were quantified by RT-qPCR (primer sequences: *Actb_F*: 5’-CGACAACGGCTCCGGCATGT-3’, *Actb_R* 5’-TCACACCCTGGTGCCTAGGGC-3’, *Igf1_F*: 5’-AGCATACCTGCCTGGGTGTCCA-3’, *Igf1*_R: 5’-TGTGTATCTTTATTGCAGGTGCGGT-3’).

### In vitro luciferase assays

Luciferase reporter constructs were generated by PCR amplification of approximately 600 bp of *Igf1* 3⍰UTR encompassing the *miR-483-3p* seed regions. These PCR products were subcloned downstream of the luciferase gene contained in the pGL3-Basic commercial vector (Promega – E1751). Mutation of the highly cross-species conserved *miR-483-3p* seed target sequence from AGGAGUG to AGGAACG (mouse) was performed using the QuickChange Site-Directed Mutagenesis kit (Agilent Technologies – 200518). The HEK-293 cells were used for *miR-483-3p* expression studies, and HepG2 cells for *miR-483-3p* knockdown studies, because of their low and high levels of endogenous *miR-483-3p* expression, respectively. Cells were transfected with 100⍰ng of reporter construct, 50⍰ng of pLacZ-Control Vector (a transfection control, Clontech – 631709) and increasing concentrations of mouse *miR-483-3p* mimic (0, 10 and 50⍰nM in HEK-293) or *miR-483-3p* 2⍰-O-Methyl antagonist (anti-human; 0, 50 and 100⍰nM in HepG2) using the Lipofectamine RNAiMAX Transfection Reagent (ThermoFisher Scientific – 13778100). Cells were harvested and the luciferase assay was performed as previously described^21^ using the Dual-Luciferase Reporter Assay System (Promega – E1910), with a transfection efficiency control (T1003, Applied Biosystems). Luminescence was detected using the GloMax Discover Microplate Reader (Promega – GM3000).

### Continuous IGF1 administration via minipumps

Human LR3-IGF1 (Preprotech – 100-11R3) was dissolved in 100 mM acetic acid in pH 7.4 sterile PBS, at a concentration of 9 μg/μl. Osmotic minipumps (Alzet – model 1004), which are designed to deliver a constant volume of 0.13 μl/hour, were pre-filled with 110 μl LR3-IGF1 or vehicle under sterile conditions. The concentration was calculated to deliver an average dose of 1.5 μg LR3-IGF1/g body weight/day. Prior to surgery, the minipumps were primed overnight in sterile PBS at 37°C. Surgery was performed on W4 *iTg*^Mir483^ mutant males, a time-point when they were above 10 g body weight. The mini pumps were implanted subcutaneously under general anaesthesia, in the subcutaneous scapular area, below the shoulder blade, just off midline. The mice were given postoperative analgesia for one day, and were monitored daily for one week. Each week, a small blood sample was collected from the tail vein and 5 μl plasma was used for mouse IGF1 measurement by ELISA and 5 μl plasma for human LR3-IGF1 measurements by liquid chromatography and massspectrometry (LC-MS). Plasma LR3-IGF1 concentrations (ng/mL) were calculated by analysis of the human LR3-IGF1-specific tryptic peptide GFYFNKPTGYGSSSR against a standard curve prepared with serial dilutions of the stock LR3-IGF1 solution in mouse plasma (50-1000 ng/mL). Plasma samples were extracted as previously described^66^ and analysed on a Waters M Class nano LC system (Milford, MA, USA), coupled to a Xevo TQ-XS triple quadrupole (Waters, MA, USA). The LR3-IGF1 specific peptide was monitored using the *m/z* transitions 556.8 / 177.0, and peptide peak area ratios were generated against a spiked internal standard with bovine insulin (Sigma – 11070-73-8).

### Statistical analyses

Statistical analyses were performed using GraphPad Prism 9 software. For two groups, statistical analyses were performed using Mann–Whitney tests or un-paired Student’s t-tests with Welch’s correction (depending on the outcome of Shapiro–Wilk tests for normal distribution). Where more than two groups were analysed, we used one-way ANOVA, followed by Tukey’s multiple comparisons tests or two-way ANOVA followed by Sidak’s corrections for multiple testing, as appropriate. For growth kinetics analyses, we used mixed-effects model (REML) tests. For all tests, *P* values⍰<⍰0.05 were considered significant.

## Acknowledgements

We thank Laura Hunter and Claire Custance (West Forvie Phenomics Center) for help with mouse husbandry; Debbie Drage, Martin George and in particular Ted Saunders (The Babraham Institute Gene Targeting Facility) for help with generating the *Igf2*^Δ(P1-P3)^ mice; James Warner and Katherine Vickers (Histopathology Core) for help with preparing tissue samples for histology; Gregory Strachan (Imaging Core) for help with confocal microscopy and cell counting/measurements using HALO; Keith Burling and Peter Barker (Core Biochemistry Assay Lab) for performing blood biochemistry measurements; Amy Warner and Sarah Grocott (Disease Model Core) for osmotic minipump implantation through surgery; Dr. Paúl Cordero for help with mRNA expression analyses; Dr. Robin Antrobus (Cambridge Institute for Medical Research Proteomics Facility) for the TMT analysis; Dr. Tim Mohun (Crick Institute, London) for help with tissue collection for HREM; Prof. Abigail Fowden for help with mouse licensing approval for part of the work on the *iTg*^Mir483^ model; Dr. Allan Bradley for providing *Mir483*^KO^ ES cells. This work was supported by the Medical Research Council (MR/J001562/1 to M.C.; MRC_MC_UU_12014/4 to M.C. and S.E.O.; MRC_MC_UU_12012/5 to the MRC Metabolic Diseases Unit; MR/M009041/1 – “Enhancing UK Clinical Research” grant supporting the TQ-XS work); the Biotechnology and Biological Sciences Research Council (BBSRC BB/F014279/2 to A.E.W. and M.B.) and the Cancer Research UK (CRUK Institute award A29252 to M.B.); N.C. was funded by the Frank Edward Elmore Fund, the Association of Physicians of Great Britain & Ireland and the Anatomical Society; Y.S. was funded by post-doctoral fellowships from the Uehara Memorial Foundation and the Japan Society for the Promotion of Science; D.C. was funded by the Erasmus placement programme; K.B.Q. was funded by Coordenação de Aperfeiçoamento de Pessoal de Nível Superior, CAPES, Brazil. Schematic representations shown in Fig. 6f–6h and Extended Data Fig. 12f were generated using BioRender (Biorender.com).

## Contributions

Conceptualization, K.S., A.E.W., M.B., S.E.O. and M.C.; Investigation: *Mir483*^KO^ (Y.S., D.C., N.C., I.Z., H.M.P., K.B.Q.), *H19*^Δ13KO^ (Y.S., D.S.F-T.), *Igf2*^KO’s^ (I.S., D.S.F-T., K.H., N.H.S.), *Mir483*^5C^ (W.N.C., I.Z., C. S.K.C., N.M.S.), *iTg*^Mir483^ (I.S., D.S.F-T., N.C., W.N.C, I.Z., K.B.Q, L.M.O., R.G.K., S.H.G., L.F.R., W.J.W.), cellular *Mir483* essays (D.F-M., M.B.); writing-original draft, M.C.; writing – review and editing: I.S., D.S.F-T., N.C., W.N.C., Y.S., I.Z., D.F-M, S.G., M.B. and M.C.; supervision, S.E.O and M.C.; funding acquisition, S.E.O. and M.C.

## Corresponding author

Correspondence to Miguel Constância

