## Supplementary data for "The imprinted *Mir483* is a growth suppressor and metabolic regulator functioning through IGF1"

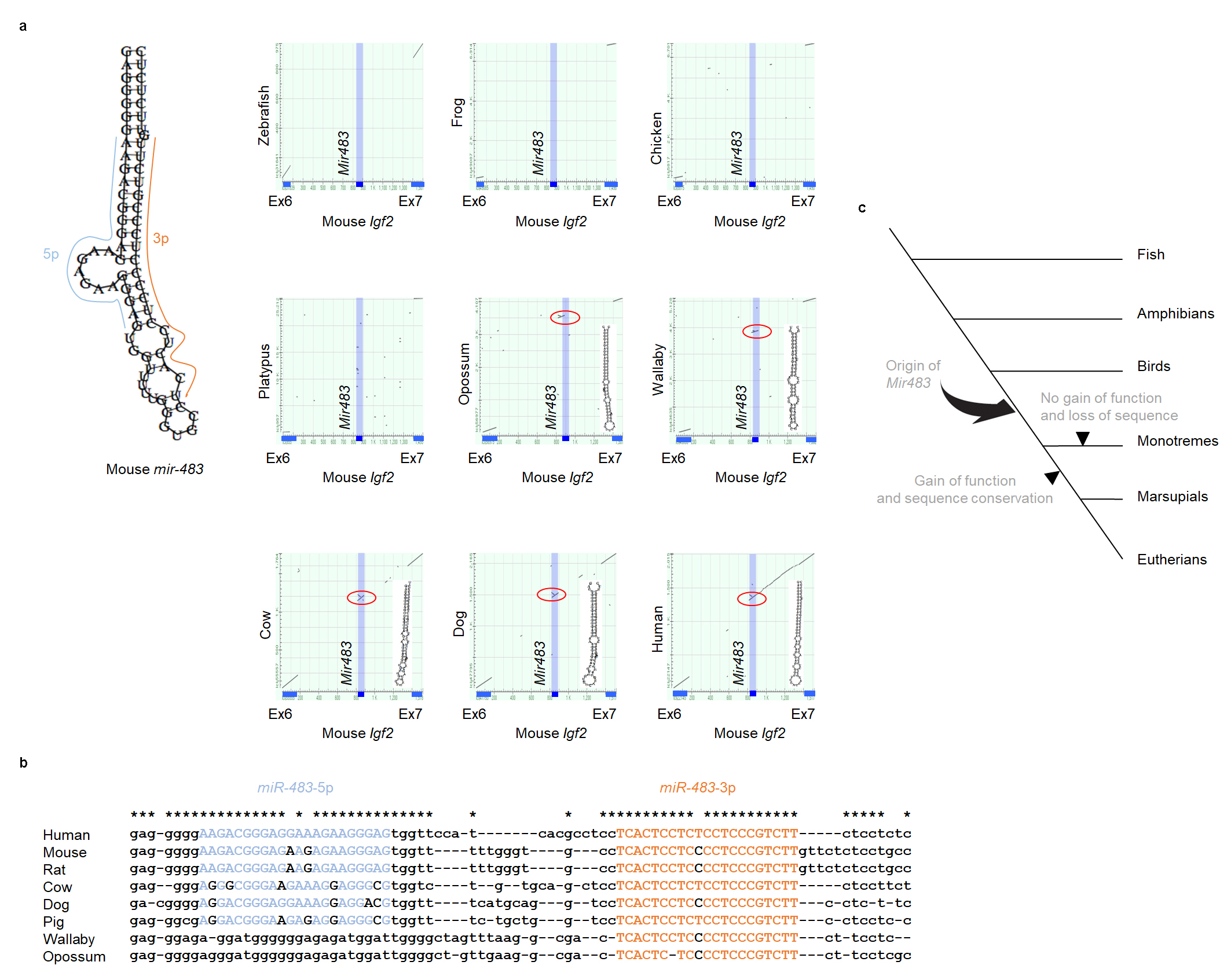

**Extended Data Fig. 1: *Mir483* sequence conservation between species.**

**a,** *Mir483* is conserved among eutherian mammals and marsupials. The Blast2 program (http://www.ncbi.nlm.nih.gov/) was used to investigate the conservation of *Mir483* sequence among vertebrates. Genomic sequences including *Igf2*’s exons 6 and 7, and intron 6 of zebrafish, frog, chicken, platypus, opossum, wallaby, cow, dog, human, and mouse were extracted from public data bases. The mouse genome was aligned with other genomes. *Igf2* exons 6 and 7 of the mouse genome and *Mir483* are highlighted by dark blue on the X axes. Grey dots indicate sequence conservation and those corresponding to *Mir483* sequence are emphasized by red circles. The secondary structures of *mir-483* in eutherian mammals and marsupials were drawn using ViennaRNA (<http://www.tbi.univie.ac.at/~ivo/RNA/>), with that of mouse being shown in the top left corner of this panel. **b,** Sequences of eutherian mammals and marsupials corresponding to *Mir483* were aligned. *miR-483-5p* and *miR-483-3p* are highlighted in blue and orange, respectively, and nucleotides conserved between human and mouse are indicated with stars. **c,** Schematic illustration of the evolution of *Mir483* in vertebrates. *Mir483* emerged in the *Igf2* intron 6 region before the divergence of the therians and monotremes. In the therian lineage, *Mir483* acquired function and became conserved in the course of evolution, whereas in the monotreme lineage, *Mir483* did not gain any function or lost it, as its sequence was not conserved.

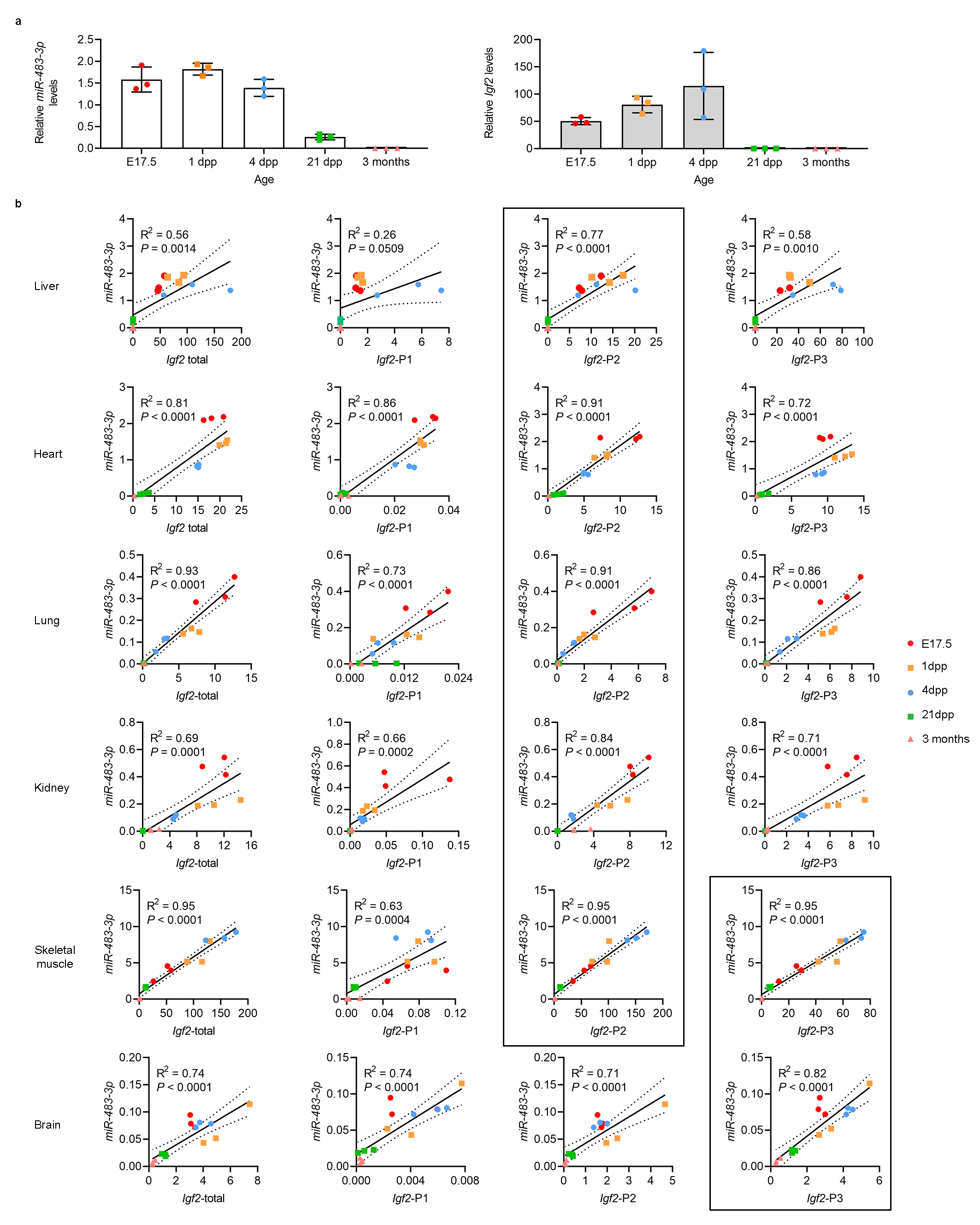

**Extended Data Fig. 2: Developmental expression of *miR-483-3p* compared to *Igf2* in prenatal and postnatal organs.**

**a,** Levels of both *miR-483-3p* and *Igf2* in the liver reach a peak in perinatal life and decrease rapidly around weaning, with very low expression levels found in adult life. **b,** Linear correlation coefficients (R^2^) between *miR-483-3p* and *Igf2* transcripts are strongest for the *Igf2*-P2 isoform in liver, heart, lung and kidney, for *Igf2*-P2 and *Igf2*-P3 in the skeletal muscle and for *Igf2*-P3 in the brain (all highlighted by black contour). For all graphs, expression of *miR-483-3p* was normalized against the geometrical mean of *Snord70*/*snoRNA234*, *Snord68*/*snoRNA202*, and *miR-26b*, and expression of *Igf2* transcript isoforms was normalized against the geometrical means of *Ppia*, *Pmm1*, *Hprt*, *Sdha*, *Tbp* and *Gapdh*. Data are presented as individual values, with averages ± SD in **a** and individual values in **b** (n=3 samples for each developmental time point).

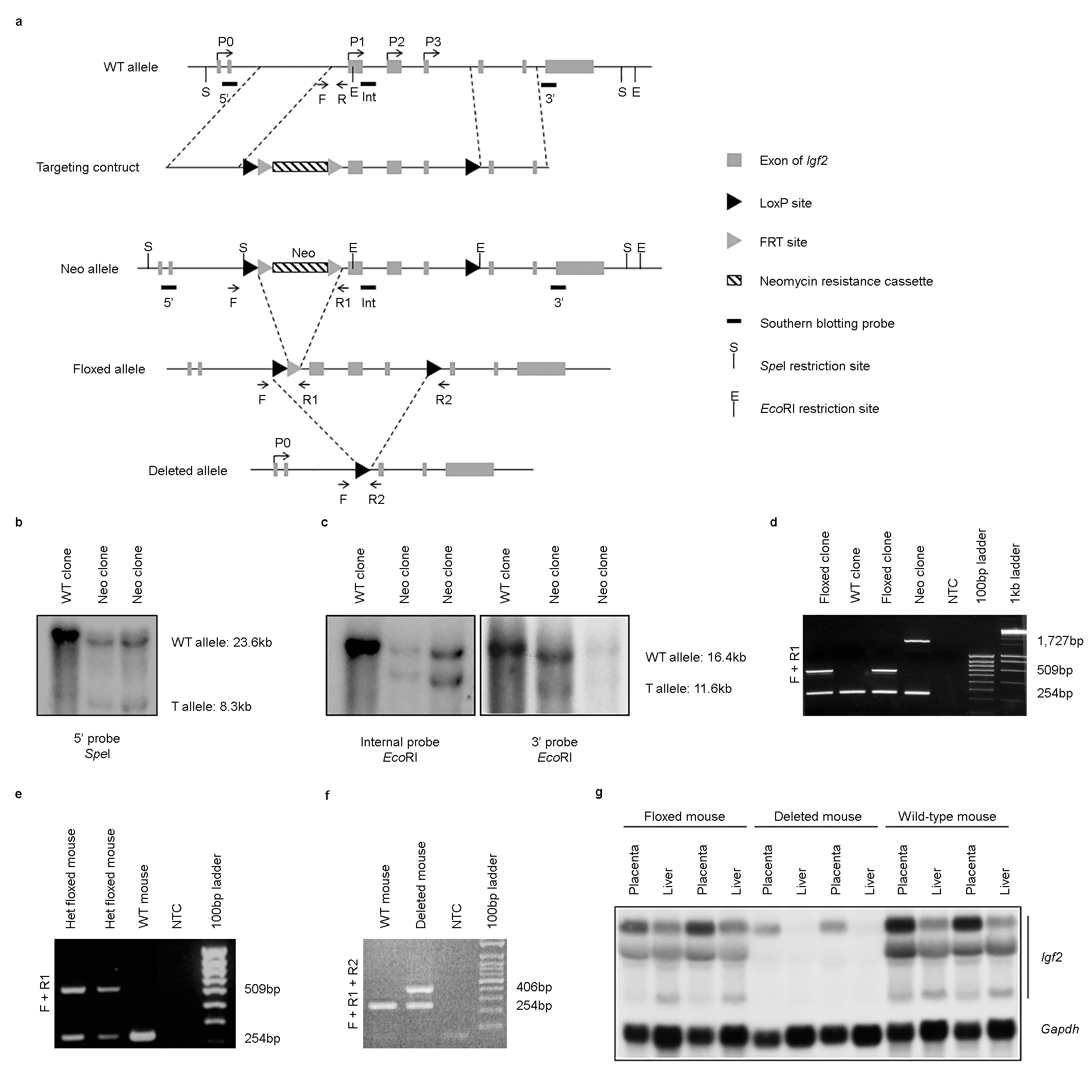

**Extended Data Fig. 3: Generation of the *Igf2*^Δ(P1-P3)^ model.**

**a,** Diagram showing the wild-type (WT) locus and the targeting construct, followed by the allele configurations after homologous recombination in ES cells, *i.e.* Neo allele, the floxed allele (after removal of the neo gene by Flpe recombinase) and the deleted allele after *Cre*-mediated recombination between LoxP sites. Probes used for Southern blot analysis (5’, Int – internal, and 3’) are shown as short horizontal black lines, selected restriction sites are indicated by E (*Eco*RI) or S (*Spe*I) and genotyping primers (F, R1 and R2) are shown by arrows. **b,** Southern blotting of DNA extracted from ES cell clones and digested with *Spe*I was used to confirm correct 5’ targeting. **c,** Southern blotting of DNA extracted from ES cell clones and digested with *Eco*RI was used to confirm correct Int and 3’ targeting. **d,** PCR confirmation of efficient removal of the neomycin cassette by FLPe-FRT recombination using primers F+R1 (NTC – no template control). **e,** PCR genotyping of heterozygous floxed mice using primers F+R1. **f,** PCR confirmation of efficient deletion of the floxed P1-P3 promoters, using primers F+R1+R2. **g,** Northern blot analysis of *Igf2* transcripts and *Gapdh* internal control in placenta and liver at E18.5 (top band observed in placentae of deleted mice corresponds to *Igf2*-P0 transcript that remains expressed upon deletion of the P1-P3 fetal *Igf2* promoters).

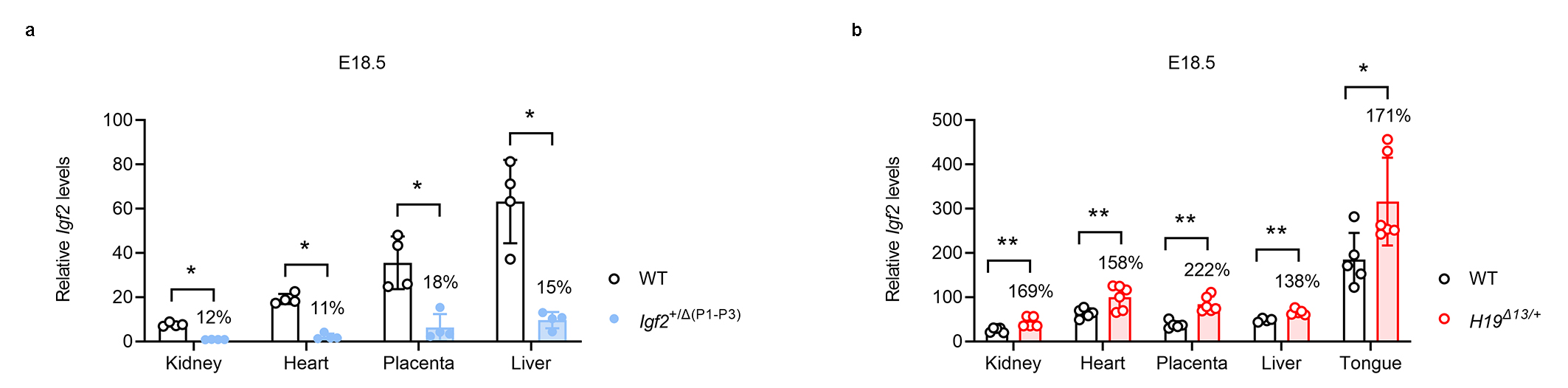

**Extended Data Fig. 4: Expression of *Igf2* in *Igf2*^+/Δ(P1-P3)^ and *H19*^Δ13/+^ models at E18.5.**

**a,** Deletion of fetal *Igf2* promoters P1-P3 from the paternal allele (*Igf2*^+/Δ(P1-P3)^) reduces drastically *Igf2* expression. **b,** Deletion of ICR1 and *H19* on the maternal allele (the *H19*^Δ13/+^ model) results in increased *Igf2* expression in multiple organs at E18.5. For both panels, *Igf2* mRNA levels were normalized against the geometrical mean of *Ppia*, *Pmm1* and *Hprt*. Data are presented as individual values with averages ± standard deviations (SD); % values above the mutant columns indicate ratios mutant/wild-type (WT); n=4-6 samples per group; * P<0.05, ** P<0.01 by Mann-Whitney tests followed by two-stage step-up (Benjamini, Krieger, and Yekutieli) FDR < 5%.

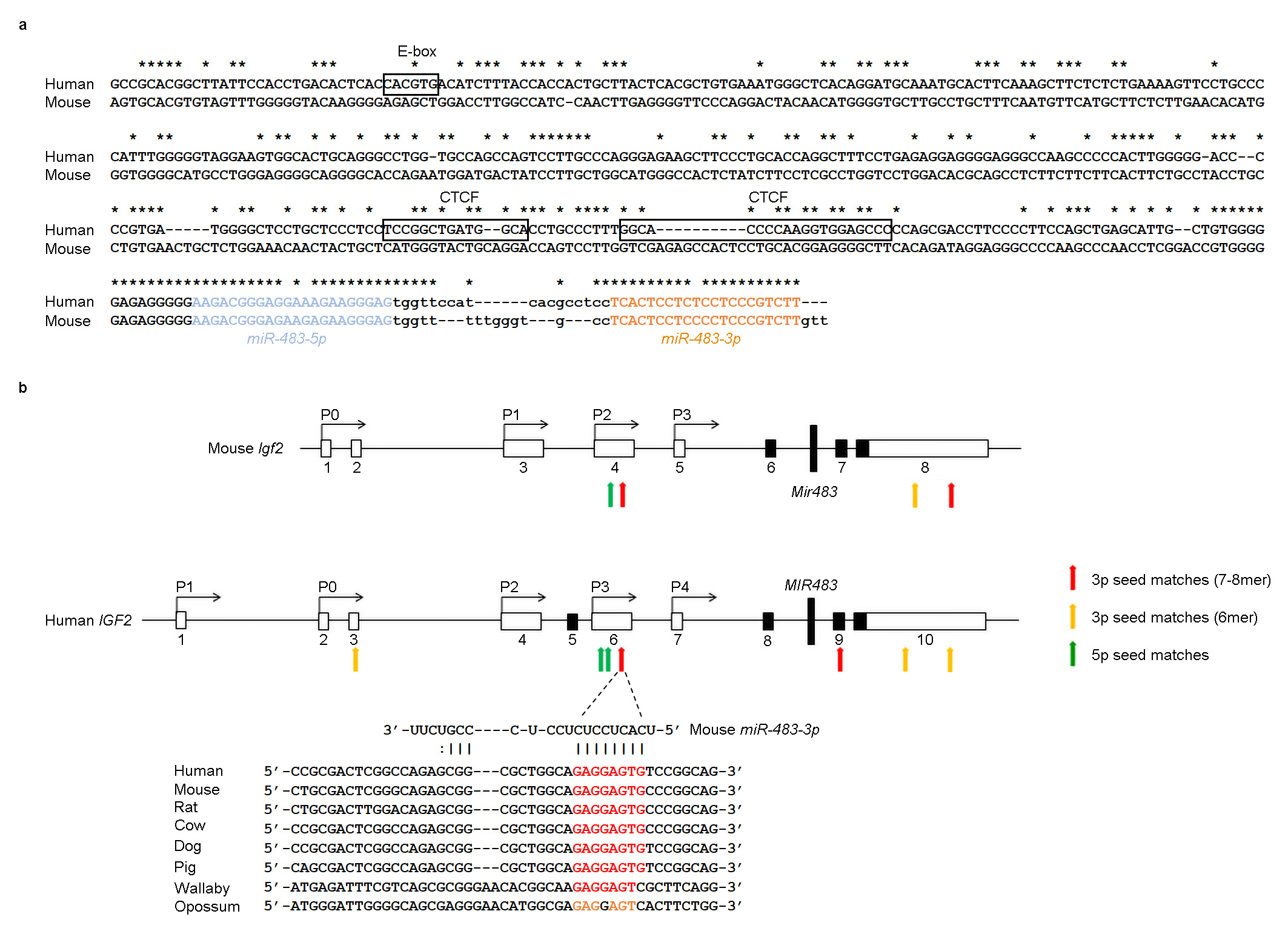

**Extended Data Fig. 5:** **Conservation between species of putative *Mir483* regulatory elements and target sequences at the *Igf2* locus.**

**a,** Sequence alignment of the human *MIR483* promoter with the equivalent sequence in the mouse shows poor conservation. The E-box and CTCF binding sites were previously identified as regulatory elements within the human *MIR483* promoter. **b,** Top: schematic representation of conserved *Igf2* 5’UTR and 3’UTR sequences between mouse and human containing *miR-483*-5p and *miR-483*-5p seed sites. Bottom: the sequence for a *miR-483*-3p seed site mapping to the untranslated mouse exon 4, driven by the *Igf2*-P2 promoter (equivalent to the untranslated human exon 6 driven by *IGF2*-P3 promoter) is conserved in the eutherian mammals and the marsupial wallaby.

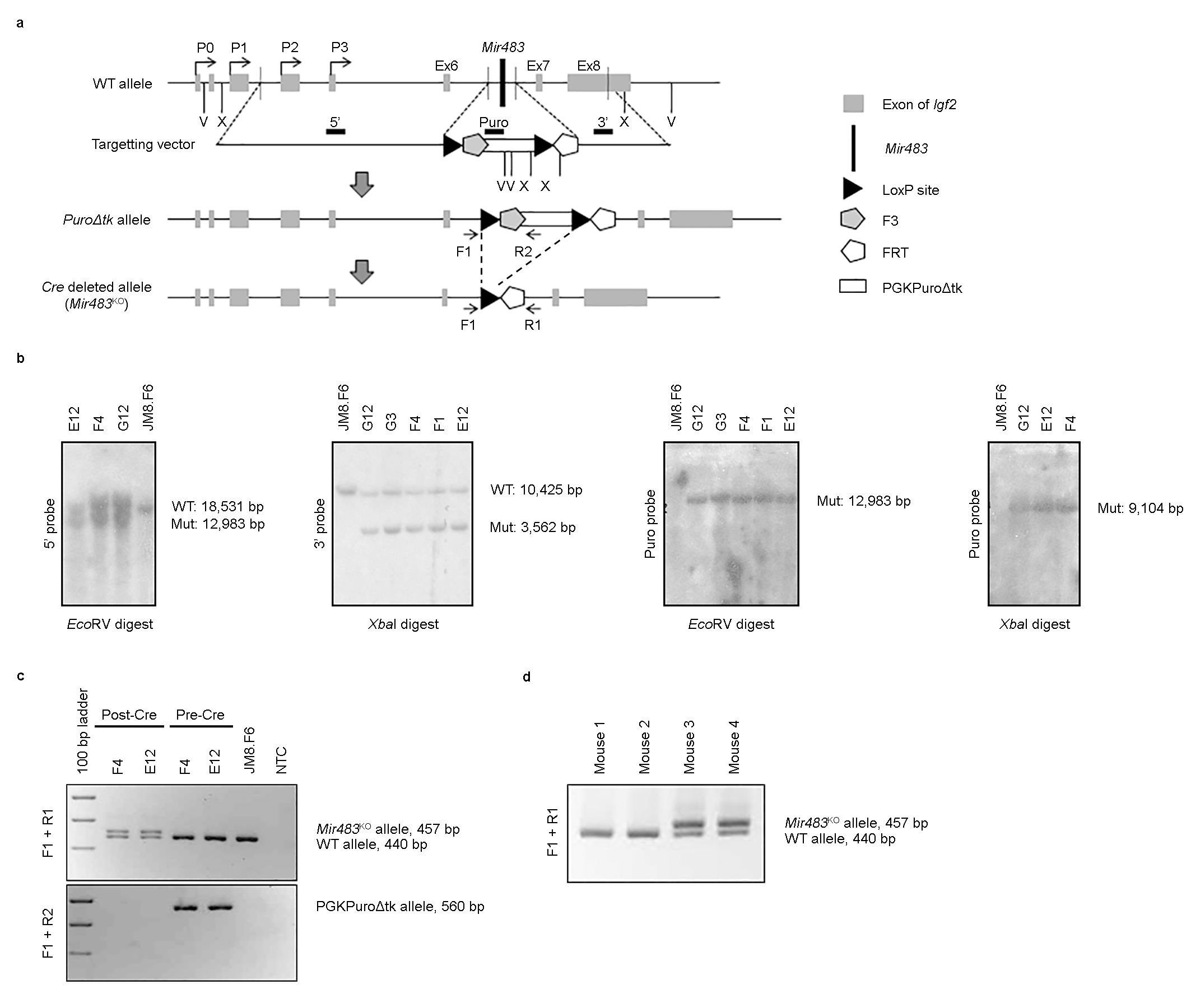

**Extended Data Fig. 6:** **Generation of the *Mir483* specific knockout.**

**a,** Diagram showing the wild-type (WT) locus and the targeting vector, the *Puro∆tk* allele after homologous recombination and the allele after *in vitro* *Cre* deletion (*Mir483*^KO^). Probes used for Southern blot analysis (5’, Puro and 3’) are shown as short black lines above the targeting vector, selected restriction sites are indicated by V (*Eco*RV) or X (*Xba*I) and genotyping primers (F1, R1 and R2) are shown by arrows. **b,** Southern blotting of ES cell clones was used to confirm correct targeting. **c,** *In vitro* *Cre* deletion in ES cell clones was confirmed by PCR (using the primers indicated in the figure) before (Pre-*Cre*) and after (Post-*Cre*) transfection with a plasmid encoding a *Cre* recombinase. Parental JM8.F6 ES cell DNA was also amplified, NTC – no template negative control. **d,** PCR genotyping in tail DNA using primers F1 and R1.

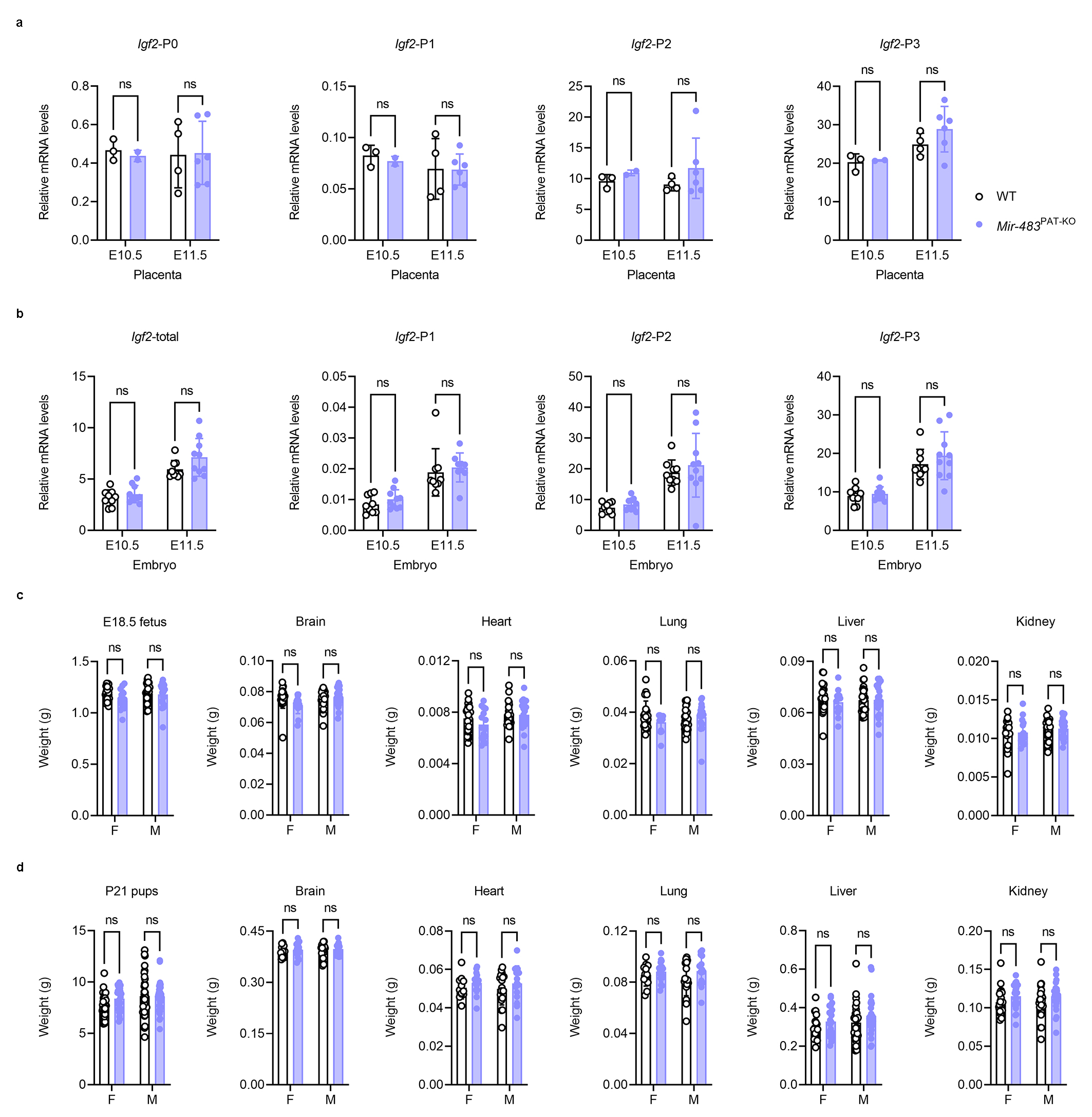

**Extended Data Fig. 7: Expression of *Igf2* isoforms and organ weights in the *Mir483*^Pat-KO^ knockout.**

**a,** Relative mRNA expression of *Igf2* isoforms at E10.5 and E11.5 measured by RT-qPCR levels in placentae (n=2-6 samples per group). Levels of *Igf2* transcripts were normalized against the geometrical mean of *Gapdh*, *Pmm1* and *Ppia*. **b,** Relative expression of total *Igf2* and its isoforms at E10.5 and E11.5 measured by RT-qPCR levels in whole embryos (n=8-10 samples per group). Levels of *Igf2* transcripts were normalized against the geometrical mean of *Gapdh*, *Pmm1* and *Ppia*. **c,** Fetus and organ weights at E18.5 (n=15-26 per group; F – females, M – males). **d,** Total body (n=33-40 per group) and organ weights (n=12-29 per group) at post-natal day 21 (P21). For all graphs, data are presented as individual values, with averages ± SD; ns – non-significant by two-way ANOVA followed by Šídák's multiple comparisons tests.

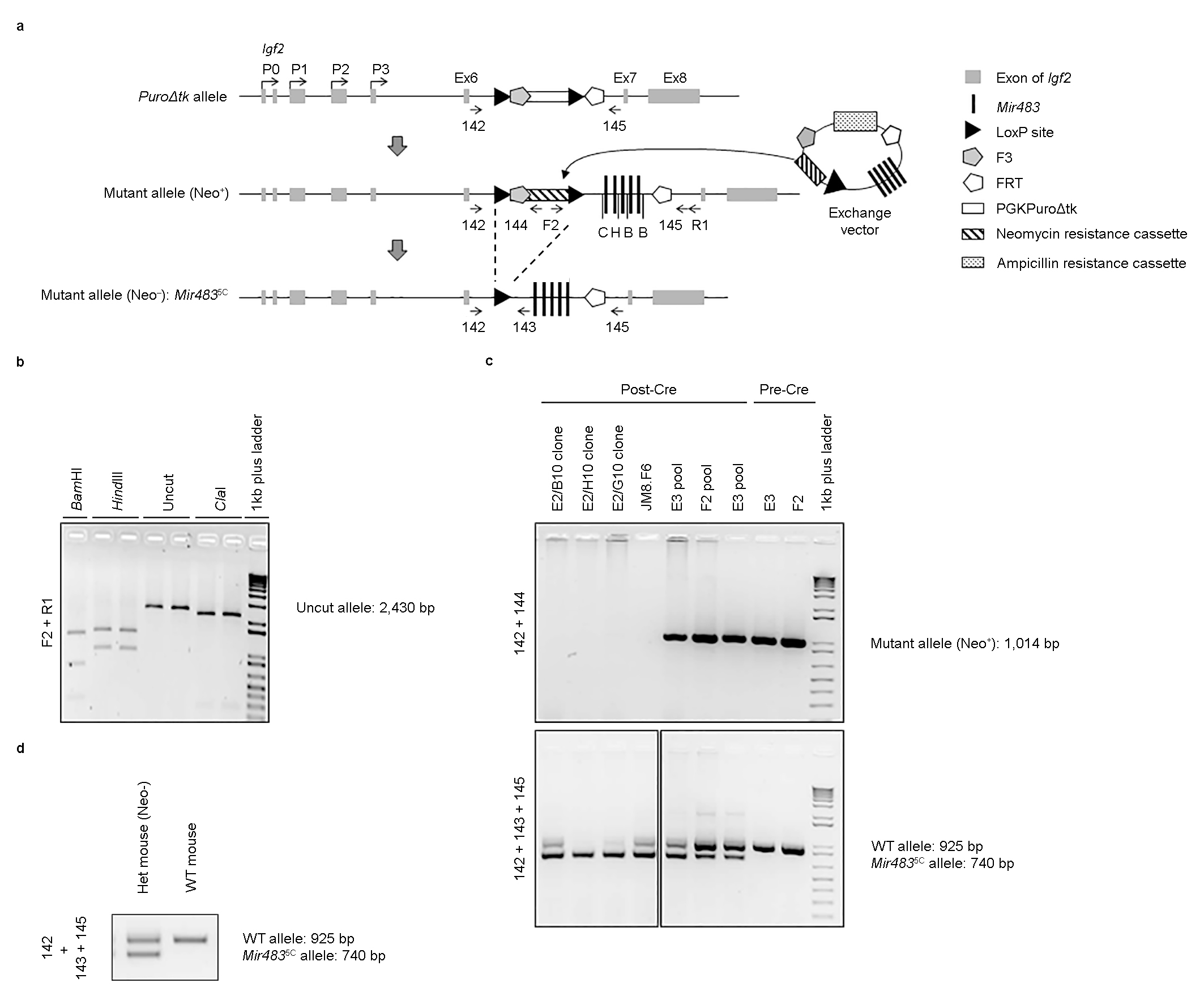

**Extended Data Fig. 8: Generation of *Mir483*^5C^ (5C) mice.**

**a,** Mice expressing 5 copies of *Mir483* from the endogenous locus were generated by replacement of the PGKPuro∆tk allele of the *Mir483*^KO^ ES cell line with a neomycin resistance cassette and 5 copies of *Mir483* using RCME (recombination mediated cassette exchange). The diagram shows the Puro∆tk allele and exchange vector, the (Neo^+^) allele after RCME, and the allele after *in vitro* *Cre* deletion (Neo^–^). Genotyping primers are depicted by arrows. Selected restriction sites are indicated by C (*Cla*I), E (*Eco*RI), H (*Hind*III) or B (*Bam*HI). **b,** PCR using primers F2 and R1 generated a 2,430 bp product indicating that RCME had occurred and the five copies of *Mir483* were present. The identity of the PCR product was confirmed by digestion with *Bam*HI, *Hind*III or *Cla*I. ***c,*** *In vitro* *Cre* deletion in ES cell clones was determined by PCR (using the primers indicated in the figure) before (Pre-Cre) and after (Post-Cre) transfection with a plasmid encoding a *Cre* recombinase. DNA pools of ESC clones and parental JM8.F6 ESC DNA were also amplified. **d,** Mice were then routinely genotyped by PCR, using primers 142, 143 and 145.

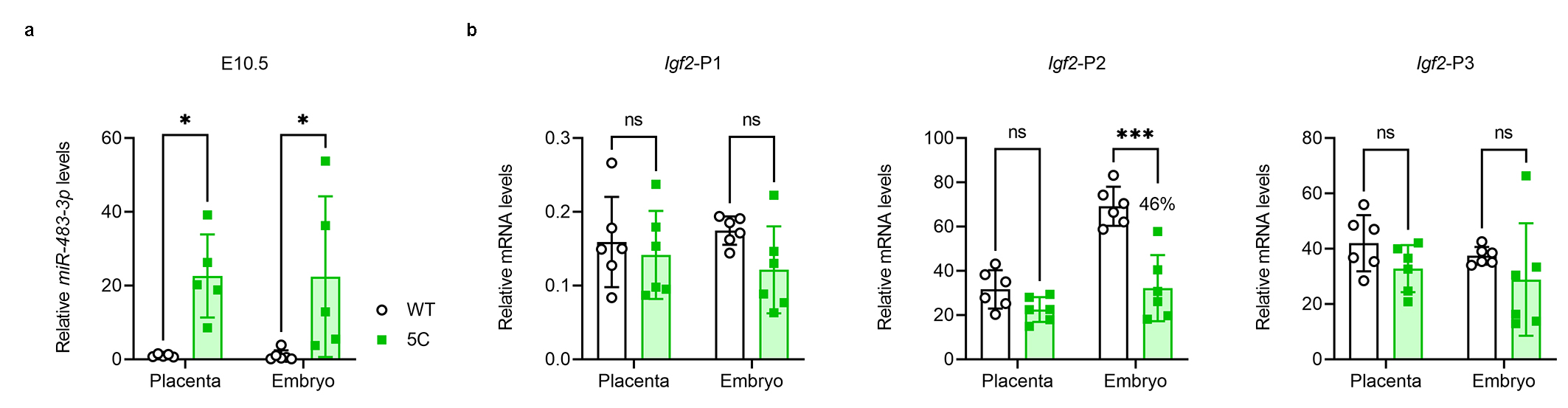

**Extended Data Fig. 9: Over-expression of *Mir483* in the *Mir483*^5C^ (5C) model is associated with down-regulation of *Igf2*-P2 isoform in embryos, but not placentae.**

**a,** Relative expression of *miR-483-3p* measured by RT-qPCR in whole embryo lysates at E10.5 (n=5-6 samples/group). Levels of *miR-483-3p* were normalized against the geometrical mean of *Snord70*/*snoRNA234*, *Snord68*/*snoRNA202*, and are presented relative to the wild-type (WT) levels, arbitrarily set to 1. **b,** Relative expression of *Igf2* isoforms at E11.5 measured by RT-qPCR levels in placenta and embryo (n=6 samples per group). Levels of *Igf2* transcripts were normalized against the geometrical mean of *Gapdh*, Sdha and *Pmm1*. For all graphs, data are presented as individual values, with averages ± SD and % indicate ratios 5C/WT; ns – non-significant, * *P*<0.05 and *** *P*<0.001 by two-way ANOVA followed by Šídák's multiple comparisons tests.

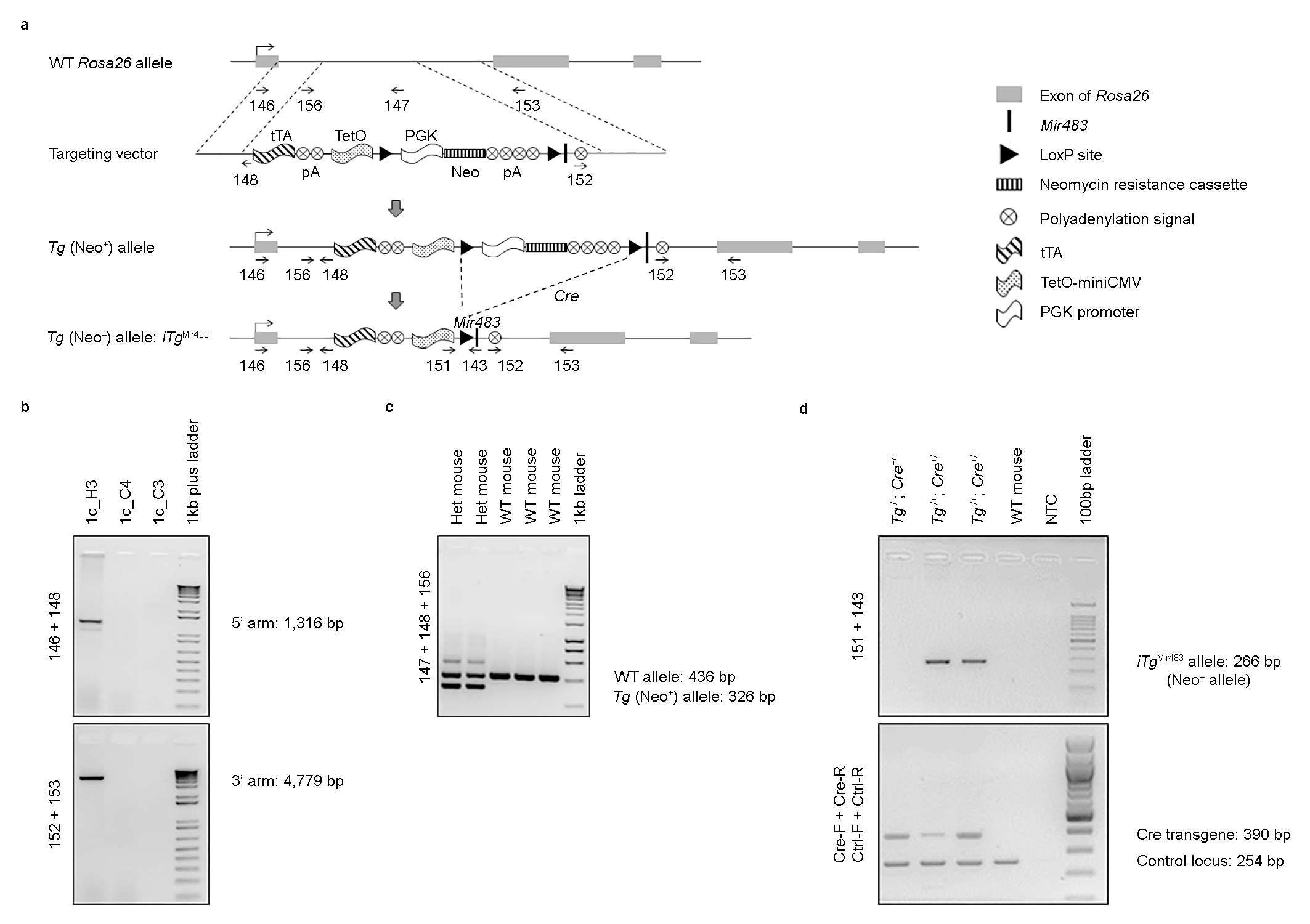

**Extended Data Fig. 10: Generation of TET-OFF inducible transgenic mice with an additional copy of *Mir483* inserted at the *Rosa26* locus (*iTg*^Mir483^).**

**a,** Diagram showing the wild-type *Rosa26* locus, the targeting vector containing a tetracycline-controlled transactivator (tTA), transgenic (Tg) (Neo^+^) allele after homologous recombination in ES cells and the Tg allele after *Cre* deletion (Neo^–^). Genotyping primers are shown by arrows. **b,** ES cell clones were screened by PCR; homologous recombination of the 5’ arm and 3’ arm was assayed using primers 146+148 and 152+153, respectively. The gel image depicts one targeted clone (1c_H3) and two non-targeted clones. **c,** Mice were then routinely genotyped using primers 147, 148 and 156. **d,** Males heterozygous for the Neo^+^ transgene were mated with female mice homozygous for a *CMV-Cre* transgene, to generate Neo^–^ progeny (*iTg*^Mir483^), which were genotyped using primers 151 and 143 (across the deleted region) and Cre-F, Cre-R, Ctrl-F and Ctrl-R (to detect the presence of *Cre*), NTC – no template negative control.

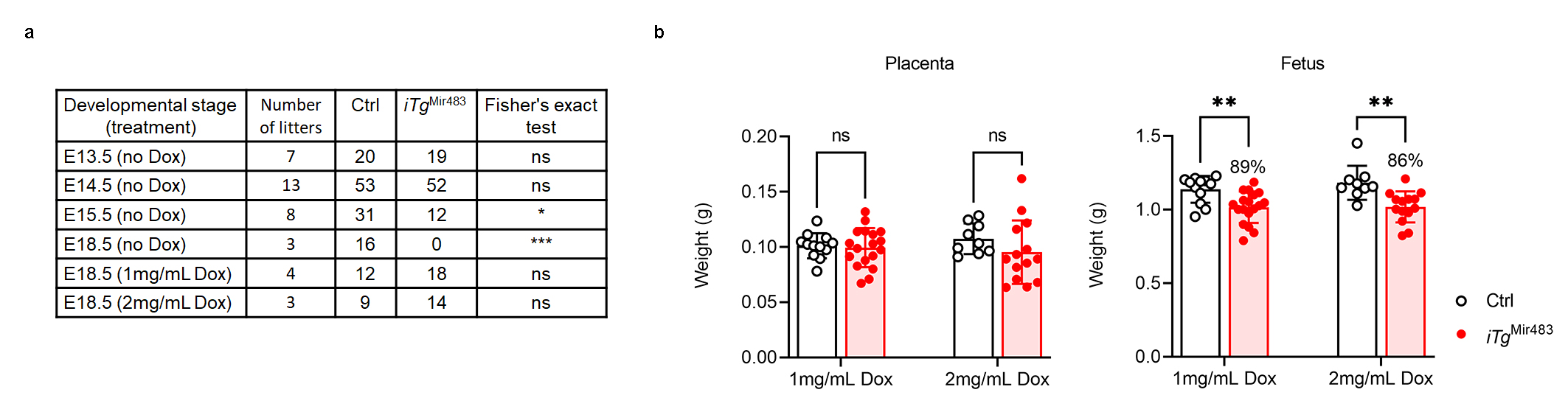

**Extended Data Fig. 11: Doxycycline administration rescues the lethality of *iTg*^Mir483^ conceptuses and partially the fetal growth restriction**

**a,** Maternal doxycycline administration in drinking water (Dox, 1mg/mL or 2 mg/mL) throughout gestation rescues the post-mid-gestation lethality of *iTg*^Mir483^ conceptuses. The table shows the distribution of live conceptuses per genotype identified at various developmental time-points during gestation. **b,** Placenta and fetal weights at E18.5 upon maternal administration of Dox (n=9-18/group). Data are presented as individual values, with averages ± SD in and % indicate ratios *iTg*^Mir483^/Ctrl; ns – non-significant, * *P*<0.05, ** *P*<0.01 and *** *P*<0.001 by Fisher’s exact tests in **a** or two-way ANOVA followed by Šídák's multiple comparisons tests in **b**.

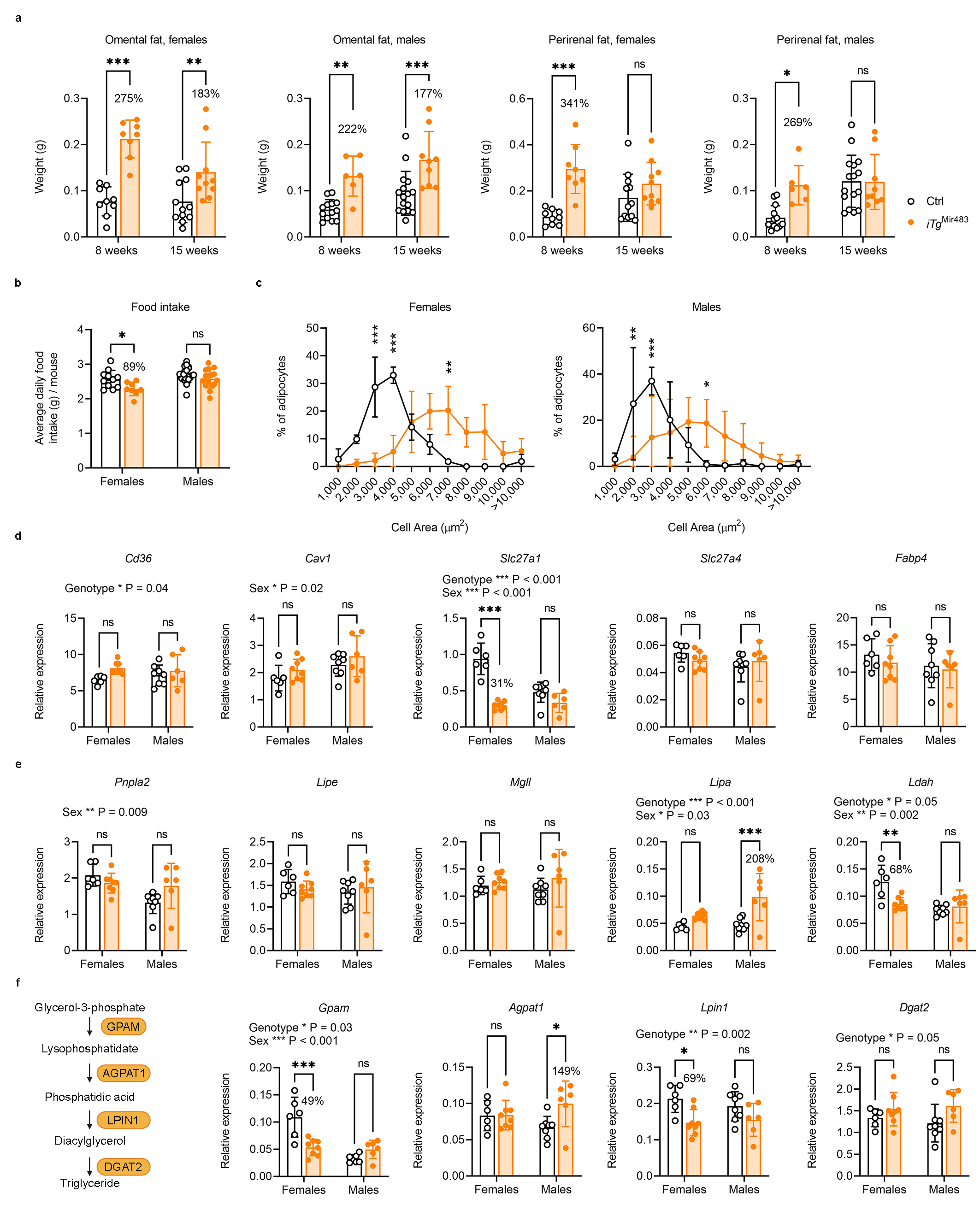

**Extended Data Fig. 12: Fat weight and morphology, and adipocyte expression patterns in the *iTg*^Mir483^ mouse model.**

**a,** Omental fat pads are significantly heavier in week 8 (W8) and W15 in *iTG*^Mir483^ adults compared to age-matched controls, while peri-renal fat pads are significantly heavier only at W8 (n=6-16 per group). **b,** Average daily food intake, measured between W3 and W4 is significantly lower in *iTG*^Mir483^ females compared to age-matched controls, with no significant differences between the two genotypes in males (n=8-20 per group). **c,** Distribution of adipocyte cell area isolated from the gonadal fat pad of W8 *iTG*^Mir483^ and age-matched controls indicates larger adipocytes in mutants of both sexes (n=2-10 per group). **d,** Expression patterns of genes encoding lipid transporters are largely unchanged in adipocytes isolated from gonadal fat of W8 *iTG*^Mir483^ and age-matched controls, with the notable expression of *Slc27a1* (also known as *Fatp1*) that is significantly down-regulated in mutant females only (n=6-8 per group). **e,** Expression patterns of genes encoding three major lipases (*Pnlpa2*, also known as *Atgl*; *Lipe*, also known as *Hsl*; *Mgll*, also known as *Mgl*) are unchanged in adipocytes isolated from gonadal fat of W8 *iTG*^Mir483^ and age-matched controls. Significant differences were observed for two minor lipases: *Lipa*, upregulated in males, and *Ldah*, down-regulated in females (n=6-8 per group). **f,** Left: diagram depicting the steps involved in the conversion of glycerol-3-phosphate (G3P) into triglycerides (TG). Right: expression patterns of genes encoding enzymes implicated in the synthesis of TG in adipocytes isolated from gonadal fat of W8 *iTG*^Mir483^ and age-matched controls (n=6-8 per group). Data are presented as individual values, with averages ± SD in **a**, **b**, **d** and **f**, or averages ± SD in **c** and % indicate ratios *iTg*^Mir483^/Ctrl; ns – non-significant, * *P*<0.05, ** *P*<0.01 and *** *P*<0.001 by two-way ANOVA followed by Šídák's multiple comparisons tests. For panels **d**, **e** and **f**, the effects of genotype and sex identified by two-way ANOVA tests are indicated above the graphs.

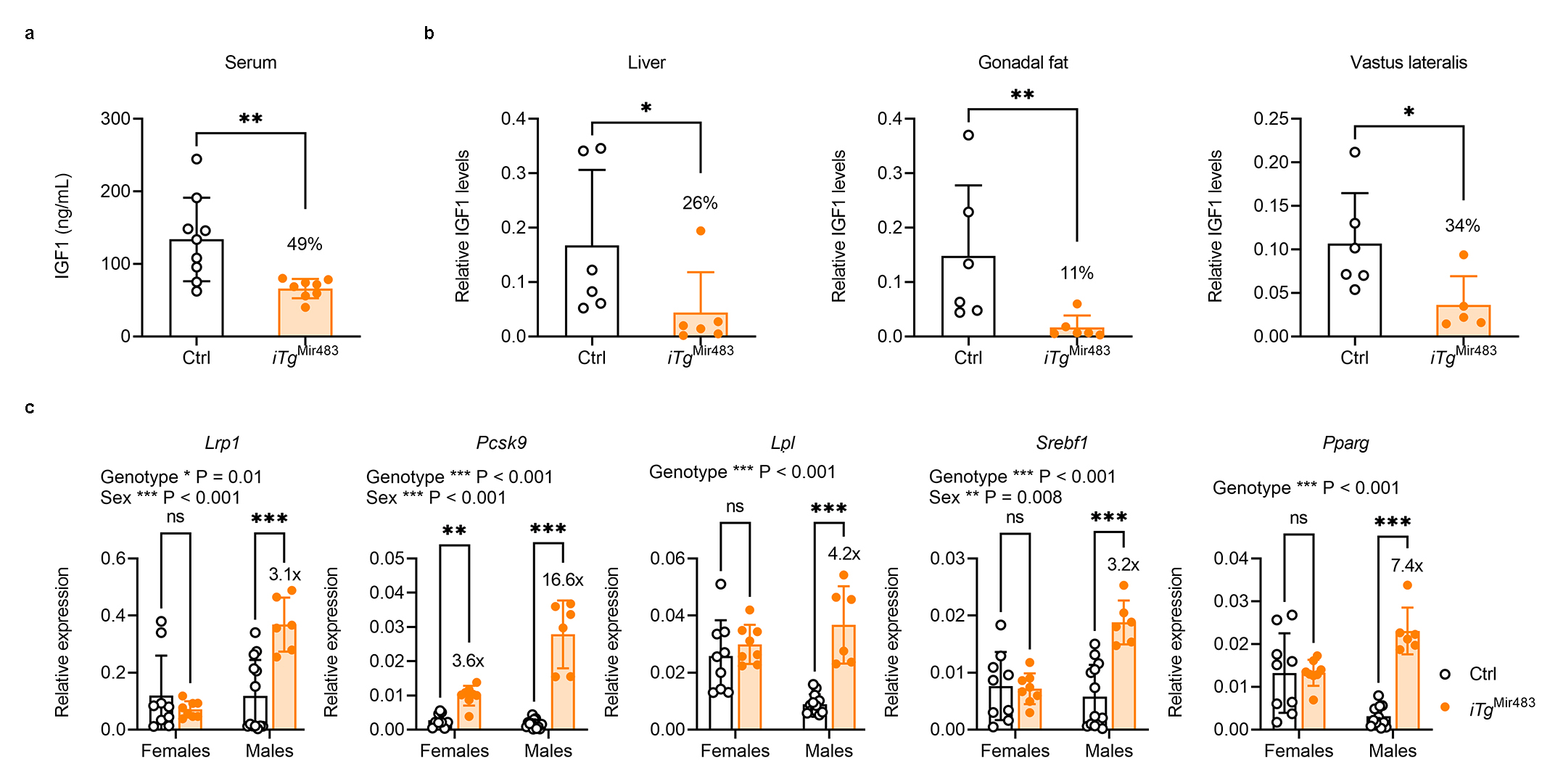

**Extended Data Fig. 13: IGF1 protein levels and mRNA levels of genes that regulate lipid trafficking or production in the liver of the *iTg*^Mir483^ mouse model.**

**a,** IGF1 protein levels measured by ELISA in serum of W3 *iTG*^Mir483^ and age-matched controls (n=8-9 per group). **b,** Relative protein levels of mature IGF1 (7.5 kDa) measured by western blot analyses in protein lysates of three organs collected from W15 males of *iTG*^Mir483^ and age-matched controls (n=5-6 per group). Levels of IGF1 were normalized against Coomassie staining in the liver and gonadal fat, and SOD1 in the *vastus lateralis*, used as internal controls for protein loading. **c,** Expression patterns of genes encoding proteins implicated in lipoprotein turnover (*Lrp1*, *Pcsk9* and *Lpl*) and key transcriptional factors that regulate lipid synthesis (*Sreb1*, *Pparg*) in livers of W8 *iTG*^Mir483^ and age-matched controls (n=6-13 per group). Data are presented as individual values, with averages ± SD and % indicate ratios *iTg*^Mir483^/Ctrl; ns – non-significant, * *P*<0.05, ** *P*<0.01 and *** *P*<0.001 by an unpaired *t*-test with Welch's correction in **a**, Mann-Whitney tests in **b** and two-way ANOVA followed by Šídák's multiple comparisons tests in **c**. For panel **c**, the effects of genotype and sex identified by two-way ANOVA tests are indicated above the graphs.

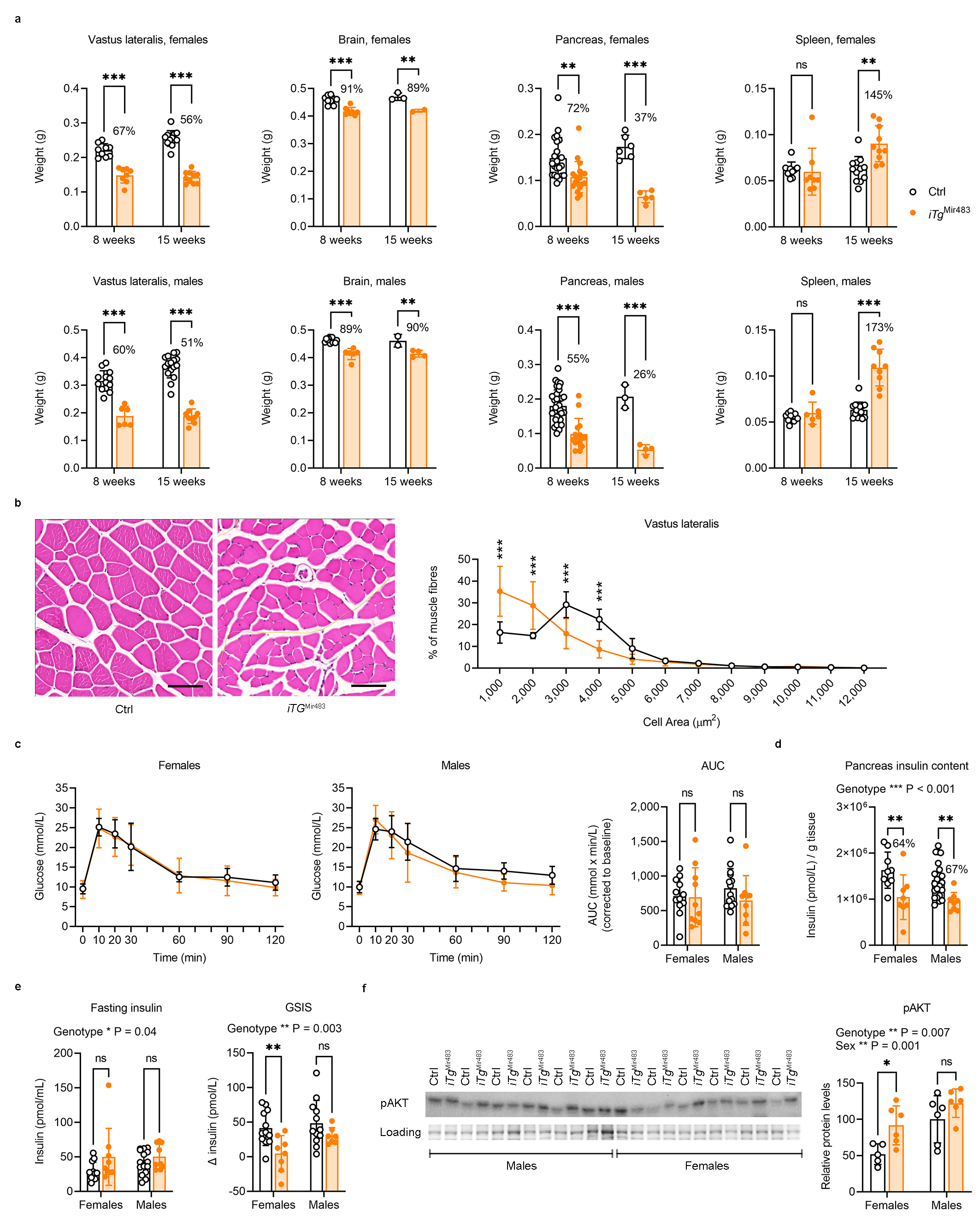

**Extended Data Fig. 14: Organ size, muscle morphology and glucose metabolism assessment in the *iTg*^Mir483^ mouse model.**

**a,** Organ weights in W8 and W15 *iTG*^Mir483^ adults compared to age-matched controls (n=2-16 per group). The *vastus lateralis* and pancreas are disproportionally smaller; the brain is proportionally smaller and the spleen is disproportionally bigger, but only at W15. **b,** The skeletal muscle fibres are smaller in the *vastus lateralis* of W15 *iTg*^Mir483^ mutant males compared to age-matched controls (representative H&E stained sections – left, and distribution of muscle fibre area – right; n=5 samples per group, scale bars are 100 µm). **c,** Glucose tolerance tests in W13 *iTg*^Mir483^ mutants and age-matched controls, with glucose administered by oral gavage (OGTTs) after six hours fasting performed in females (n=10-12/genotype) and males (n=9-15/genotype). First two panels show changes in blood glucose concentrations (y-axis), from basal pre-treatment values, with time (x-axis), after glucose administration. The graph on the far right shows area under curve (AUC) calculated during OGTTs using the trapezoid rule and normalised to basal glucose levels. **d,** Total pancreas insulin content in W18 *iTg*^Mir483^ mutants and age-matched controls after overnight fasting (n=8-22 per group). **e,** Left: insulin levels measured in plasma in W13 *iTg*^Mir483^ mutants and age-matched controls, after six hours fasting and prior to the start of the OGTT (n=9-16 samples per group). Right: glucose-stimulated insulin secretion (GSIS) measured in plasma at minute 20 during OGTT (n=7-12 samples per group). **f,** pAKT levels normalized to protein loading (as assessed following Coomasie R-250 dye staining) in the gonadal fat of W8 *iTG*^Mir483^ adults compared to age-matched controls (left – western blotting, right – quantification; n=6 per group). Data are presented as individual values, with averages ± SD in **a**, **c** (far right), **d**, **e** and **f** (right) or as averages ± SD in **b** and **c** (left side), and % indicate ratios *iTg*^Mir483^/Ctrl; ns – non-significant, ** *P*<0.01 and *** *P*<0.001 by two-way ANOVA followed by Šídák's multiple comparisons tests in **a**, **b**, **c**, **d**, **e** and **f**. For panels **d**, **e** and **f**, the effects of genotype or sex identified by two-way ANOVA tests are indicated above the graphs.

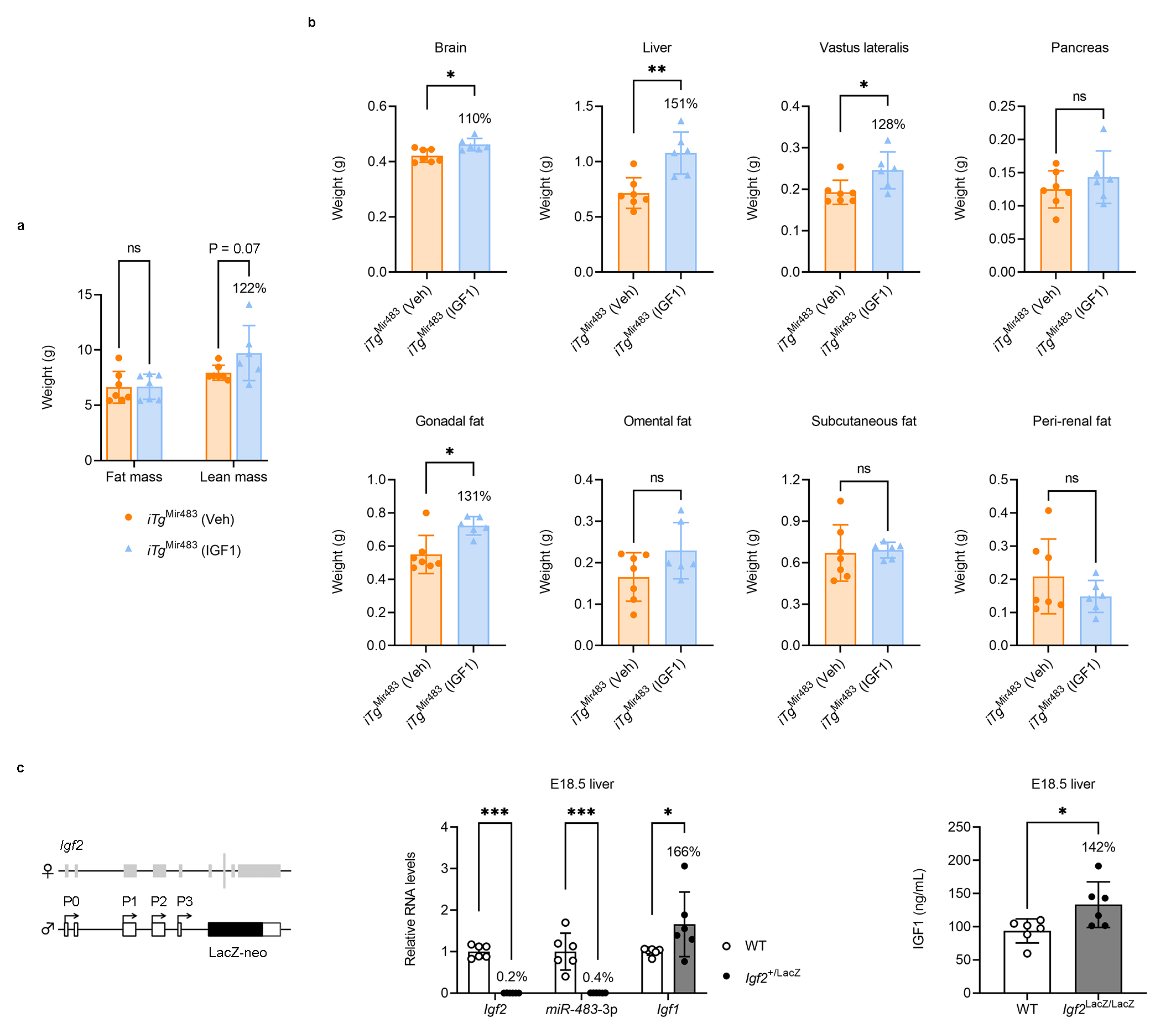

**Extended Data Fig. 15: Impact of IGF1 infusion on body composition in the *iTg*^Mir483^ mouse model and additional *in vivo* evidence for *Igf1* as target of *miR-483*.**

**a,** Body composition measured by TD-NMR in W8 *iTg*^Mir483^ male mice exposed to vehicle or IGF1 via minipumps (n=6-7 per group). **b,** Organ weights in W8 *iTG*^Mir483^ male mice exposed to vehicle or IGF1 via minipumps (n=6-7 per group). **c,** Left: schematic representation of the *Igf2*^+/LacZ^ model in which the coding exons 4-6 of *Igf2* and *Mir483* are replaced by a LacZ cassette. Middle: relative RNA levels for *Igf2*, *miR-483-3p* and *Igf1* in the liver of E18.5 *Igf2*^+/LacZ^ mutants and WT littermate controls (data was normalized against the geometrical means of *Ppia* and *Gapdh* for *Igf2* and *Igf1*, and against *Snord70*/*snoRNA234* for *miR-483*-3p; n=6 samples per group). Right: measurement of IGF1 protein by ELISA in the liver of E18.5 *Igf2*^LacZ/LacZ^ mutants and WT littermate controls (n=6 per group). Data are presented as individual values with averages ± SD and % values indicate ratios *iTG*^Mir483^ (IGF1)/ *iTG*^Mir483^ (Veh) in **a** and **b** or *Igf2*^+/LacZ^/WT or *Igf2*^LacZ/LacZ^/WT in **c**; ns – non-significant, * *P*<0.05, ** *P*<0.01 and *** *P*<0.001 by two-way ANOVA followed by Šídák's multiple comparisons tests in **a** and **c** (middle), or Mann-Whitney tests in **b** and **c** (right).

**Supplementary Table 1. List of proteins detected by TMT (Tandem Mass Tag) in E10.5 embryos in the *Mir483*^5C^ model.**

**Supplementary Table 2. Overview of developmental defects observed by HREM (High-Resolution Episcopic Microscopy) in *iTg*^Mir483^ embryos at E14.5 (n=6).**

| **Organ/system** | **Specific defect** | **Frequency** |
| --- | --- | --- |
| Cardio-vascular | Atrium and AV junction defects | 100% |
|  | Double outlet right ventricle (DORV) with associated ventricle septum defect |  |
|  | Malformations of intrathoracic arteries (interruption of aortic arch, aortic coarctation, right sided aortic arch, left sided lusoria artery, connections between subclavian artery and pulmonary trunk) |  |
|  | Abnormalities of head arteries |  |
|  | Abnormalities of the ductus venosus and portal vein |  |
| Urogenital tract | Abnormal remodeling of the metanephrotic tissue and its ducts | 83% |
|  | Abnormalities of kidneys or ureters (abnormal pelvis renalis to absent ureter and additional ureteral buds) |  |
|  | Malformations of the Wolff and Müller ducts |  |
| Skeleton | Absent or abnormal acromion with abnormal acromioclavicular joints | 83% |
|  | Abnormal otic vesicles |  |
|  | Thoracoschisis |  |
|  | Abnormal nasal cavities and associated head bones |  |
|  | Abnormal tail morphology |  |
| Thymus | Abnormal topology of the thymus | 67% |
| Liver/bile ducts | Enlarged sinusoidal spaces | 67% |
|  | Additional ducts branching from the cystic duct to enter the liver |  |
| Eye and eye muscle | Eyes with missing lenses | 67% |
|  | Eye muscle abnormalities |  |
|  | Retro-lental blood |  |
| Nervous system | Brain defects | 50% |
|  | Holoprosencephaly |  |
|  | Smaller superior cervical ganglion |  |
| Thyroid | Unilateral absence of the thyroid gland lobe | 33% |

**Supplementary Table 3. Primers used for genotyping mouse strains by PCR.**

| Mouse strain | Primer | Sequence (5’-3’) | Primer | Sequence (5’-3’) | Amplicon (bp) |
| --- | --- | --- | --- | --- | --- |
| *Igf2*^Δ(P1-P3)^ | F | ATGTCTCCAATCCTTGAACACTG | R1  R2 | GCAGTGGGAGAAATCAGAACC  GCTTTTTTAGTGGTGGGAGGC | WT – 254  Floxed – 509  Del – 406 |
| *H19*^Δ13^ | F | TGCCACAGAGGAA-GAAACCAG | R1  R2 | AGTCATAGCCGAATAGCC  TTCAGTCACTTCCCTCAGCCTC | Δ13 – 895  WT – 494 |
| *Mir483*^KO^ | F1 | TACCTGCCTGTGAACTGCTCTG | R1 | ATCTGGTGCCTCCTGTCTGGTA | WT – 440  KO – 457 |
| *Mir483*^5C^ | 142 | CACGCTTCAGTTTGTCTGTTCG | 143  145 | AAGAATCGATACCGTCGACCTC  CTGGAGTGGTTTGGAAAACAGG | WT – 925  5C – 740 |
| *iTg*^Mir483^ | 156  151 | TCCCAAAGTCGCTCTGAGTT  AGGGAGTGGTAAACTCGACC | 147  148  143 | GGCGGATCACAAGCAATAAT  GAAAGACCGCGAAGAGTTTG  AAGAATCGATACCGTCGACCTC | WT – 436  Neo^+^ – 326  Neo^–^ – 266 |
| *CMV*-Cre | Cre-F  Ctrl-F | CGAGTGATGAGGTTCGCAAG  ATGTCTCCAATCCTTGAACACTG | Cre-R  Ctrl-R | TGAGTGAACGAACCTGGTCG  GCAGTGGGAGAAATCAGAACC | Cre – 340  WT – 254 |
| *Igf2*^KO^ | F | TTACAGTTCAAAGCCACCACG | R1  R2 | GCCAAAGAGATGAGAAGCACC  GCCAAACACAGTAAAAAGAAATGC | WT – 324  Floxed – 449  Del – 384 |
| *Igf2*^LacZ^ | F | TCCTCAAGGGTTTCTTACAGTTC | R1  R2 | CCTCGACTAAACACATGTAAAGC  GACAAACTGAAGCGTGTCAAC | WT – 420  LacZ – 807 |

**Supplementary Table 4. Primers and TaqMan probes used for RT-qPCR.**

| Gene | Primer | Sequence (5’-3’) | Primer | Sequence (5’-3’) | Amplicon (bp) |
| --- | --- | --- | --- | --- | --- |
| *miR-483-3p* | mmu481853_mir (ThermoFisher Scientific #A25576) | | | | |
| *miR-483-5p* | mmu481180_mir (ThermoFisher Scientific #A25576) | | | | |
| *Snord70/* *snoRNA234* | 001234 (ThermoFisher Scientific #4427975) | | | | |
| *Snord68/* *snoRNA202* | 001232 (ThermoFisher Scientific #4427975) | | | | |
| *miR-26b* | mmu481662_mir (ThermoFisher Scientific # A25576) | | | | |
| *Igf2* | F | AGTCCGAGAGGGACGTGTCTA | R | CGGACTGTCTCCAGGTGTCAT | 102 |
| *Igf2*-P0 | F | GAGGAAGCTCTGCTGTTTGG | R | CAAAGAGATGAGAAGCACCAAC | 92 |
| *Igf2*-P1 | F | GACAAGGGTCTGACTTGGGA | R | CGTAGGAGAAGTGACGAGGC | 113 |
| *Igf2*-P2 | F | GTCGACCCTAACCGAGCTG | R | AAGCAGAGGAGAGGATGCAA | 103 |
| *Igf2*-P3 | F | TGGACATTAGCTTCTCCTGTGA | R | GCTGGAAGAGGATGAAGACAG | 61 |
| *Igf1* | F | GCTGGTGGATGCTCTTCAGTT | R | CTCATCCACAATGCCTGTCTG | 112 |
| *Ppia* | F | AAGGGTTCCTCCTTTCACAGAA | R | GATGCCAGGACCTGTATGCTT | 146 |
| *Pmm1* | F | ATCCGGGAGAAGTTTGTGGAA | R | GCTGTCTTCATCCAGGCTGTC | 144 |
| *Hprt* | F | CATTATGCCGAGGATTTGGAA | R | CCTTCATGACATCTCGAGCAA | 88 |
| *Tbp* | F | AACAACAGCCTTCCACCTTATG | R | TGTTCTGAATAGGCTGTGGAGT | 127 |
| *Gapdh* | F | ACAACTCACTCAAGATTGTCAGCA | R | ATGGCATGGACTGTGGTCAT | 121 |
| *Actb* | F | GATCAAGATCATTGCTCCTCCTG | R | AGGGTGTAAAACGCAGCTCA | 183 |
| *Acaca* | F | ACGTGCAATCCGATTTGTTGT | R | CCAGCCCACACTGCTTGTA | 180 |
| *Fasn* | F | TGCACCTCACAGGCATCAAT | R | GTCCCACTTGATGTGAGGGG | 104 |
| *Elovl6* | F | ACCCGAACTAGGTGACACGA | R | AGTCATGAACCAACCACCCC | 142 |
| *Scd1* | F | TGGTGAACAGTGCCGCGCAT | R | CGGCACCCAGGGAAACCAGG | 90 |
| *Acat2* | F | TACCTCAGTCGCAGACAGGA | R | TGAAGGAGCCTATAGCGGTG | 119 |
| *Hmgcs1* | F | CTGCTATTCTGTCTACCGCAAA | R | GGAACATCCGAGCTAGAGATTTC | 146 |
| *Hmgrc* | F | GGAATGCCTTGTGATTGGAGTT | R | CTCTAGGACCAGCGACACAC | 142 |
| *Abca1* | F | AGGGCATGTGGGAAGAACTC | R | TGTTCCCAAAACTGGTCATTGC | 115 |
| *Gpam* | F | TGCAACACTGAAATGGAAGGAG | R | ATAACATTCCGCAAACCCAGAG | 138 |
| *Agpat1* | F | AGCTCCAGTGCCAAGTATTTCT | R | GTATTTGACGTGGAGCAGCAG | 150 |
| *Lpin1* | F | GGAGACAACGGAGAAGCATTTT | R | GTTCCTCTTCAGCTGGCTTTC | 129 |
| *Lpin2* | F | CCTGAGGTCCAAGGAGAAAGT | R | GCTTCCCCATTATCACCCAATT | 87 |
| *Dgat2* | F | GCTGCAGGTCATCTCAGTACTA | R | TGCAGAAGGTGTACATGAGGAT | 89 |
| *Cd36* | F | TGCTGGAGCTGTTATTGGTG | R | GGTGCCTGTTTTAACCCAGTT | 148 |
| *Cav1* | F | ATACGTAGACTCCGAGGGACA | R | ACGTCGTCGTTGAGATGCTT | 174 |
| *Slc27a1* | F | TTCTCGTGGGCCAGATCAAC | R | AGCACGTCACCTGAGAGGTA | 136 |
| *Slc27a4* | F | CCAAAGCTGCCATTGTGGTG | R | ATCCCCACGATGTTTCCTGC | 136 |
| *Fabp4* | F | AGCTGGTGGTGGAATGTGTTAT | R | CCTCTTCCTTTGGCTCATGC | 76 |
| *Pnpla2* | F | GGAGGAATGGCCTACTGAACC | R | ATCCTCTTCCTGGGGGACAA | 71 |
| *Lipe* | F | CAAGCCCCAAAAGACCACATC | R | CTTCTTCAAGGTATCTGTGCCC | 127 |
| *Mgll* | F | AGACGGACAGTACCTCTTTTGT | R | ATGTCCAGCCCCTTCAACATAT | 135 |
| *Lipa* | F | AGATAATCATGCGCTGGGGATA | R | AAGATACACAACTGGTCTGGGA | 137 |
| *Ldah* | F | GTAGGCACCTATATGACCCTTC | R | CGATAGTTGGGAAGAGCAGAAA | 85 |
| *Lrp1* | F | GCCCACGTGCTACTGTAACA | R | ATGTGAAGGAGCCATCTGTGTT | 128 |
| *Pcsk9* | F | CTTGGTGAAGATGAGCAGTGAC | R | GGATAATTCGCTCCAGGTTCC | 122 |
| *Lpl* | F | TGTGAAATGCCATGACAAGTCT | R | CACTTTCAAACACCCAAACAAGG | 150 |
| *Srebf1* | F | ACTTTTCCTTAACGTGGGCCT | R | AGCTGGAGCATGTCTTCGAT | 153 |
| *Pparg* | F | TTTAAAAACAAGACTACCCTTTACTGAAATT | R | AGAGGTCCACAGAGCTGATTCC | 95 |
| *Ttc36* | F | CCTTTGGAGATGTTGTTGGATT | R | CAACTGTGCTTGAGGGAAAACT | 80 |
| *Arhgdig* | F | AGATTGTCAGTGGCCTCAAATG | R | AATTCATACTCCTGGGCTCTGG | 112 |
| *Lep* | F | CCCAAAATGTGCTGCAGATAG | R | CCAGCAGATGGAGGAGGTC | 63 |

**Supplementary Table 5. Primary and secondary antibodies used in Western blotting experiments.**

| Antibody | Dilution | Supplier |
| --- | --- | --- |
| Rabbit anti-phospho-AKT (Ser 473) | 1:1,000 | Cell Signaling Technology (#9271) |
| Polyclonal Goat anti-Mouse IGF-II | 1:2,000 | R&D Systems (#AF792) |
| Polyclonal Goat anti-Mouse IGF-I | 1:2,000 | R&D Systems (#AF791) |
| Rabbit polyclonal anti-SOD1 | 1:1,000 | Abcam (#ab183881) |
| Goat anti-Rabbit IgG (HRP) | 1:20,000 | Abcam (#ab6721) |
| Rabbit anti-Goat IgG (HRP) | 1:10,000 | ThermoFisher Scientific (#31433) |
